# Spatial confinement of *Trypanosoma brucei* in microfluidic traps provides a new tool to study free swimming parasites

**DOI:** 10.1101/2023.07.03.547584

**Authors:** Mariana De Niz, Emmanuel Frachon, Samy Gobaa, Philippe Bastin

**Author notes:** Equally contributed (Samy Gobaa, Philippe Bastin).

## Abstract

*Trypanosoma brucei* is the causative agent of African trypanosomiasis and is transmitted by the tsetse fly (*Glossina spp.*). All stages of this extracellular parasite possess a single flagellum that is attached to the cell body and confers a high degree of motility. While several stages are amenable to culture *in vitro,* longitudinal high-resolution imaging of free-swimming parasites has been challenging, mostly due to the rapid flagellar beating that permanently twists the cell body. Here, using microfabrication, we generated various microfluidic devices with traps of different geometrical properties. Investigation of trap topology allowed us to define the one most suitable for single *T. brucei* confinement within the field of view of an inverted microscope while allowing the parasite to remain motile. Chips populated with V-shaped traps allowed us to investigate various phenomena in cultured procyclic stage wild-type parasites, and to compare them with parasites whose motility was altered upon knockdown of a paraflagellar rod component. Among the properties that we investigated were trap invasion, exploratory behaviour, and the visualization of organelles labelled with fluorescent dyes. We envisage that this “Tryp-Chip” will be a useful tool for the scientific community, as it could allow high-throughput, high-temporal and high-spatial resolution imaging of free-swimming *T. brucei* parasites.

## Introduction

*Trypanosoma brucei* is a kinetoplastid parasite causative of African Trypanosomiasis, or sleeping sickness, in humans, and nagana in cattle. Parasites are transmitted by the bite of infected tsetse flies (*Glossina spp.*). These vectors are restricted to sub-Saharan Africa. Despite the low prevalence of infections in humans at present (with under 700 cases reported to the World Health Organization in 2020) [1], African trypanosomiasis continues to be a neglected tropical disease of significant public health relevance. This is because despite being close to eradication, the final steps to achieve it include many political, public health, economic, and logistical complex challenges so far difficult to overcome [2]. Moreover, nagana continues to be of major veterinary importance. Beyond its clinical relevance, *T. brucei* is an interesting model organism for studying micro-swimmer motility, as it has a unique flagellar anatomy, and a peculiar motility type with the flagellum leading at the anterior end. Trypanosomes are efficient swimmers both *in vitro* and *in vivo*. Their motility results from the beating of their flagellum, which has been a subject of research for decades [3,4]. Structurally, the *T. brucei* flagellum wraps around the cell body in a left-handed helix. While early research suggested that this structure causes the entire cell to rotate with uniform handedness as the flagellum beats, high-speed imaging has since revealed that the *T. brucei* cell body does not just continuously rotate with uniform helical handedness [5,6]. Instead, the handedness of the helical flagellum changes simultaneously with changing the direction of helix rotation. Cell propulsion is ultimately driven by a bihelical waveform, in which helical waves of alternating handedness propagate along the flagellum and are separated by a topological feature described as a ‘kink’. This results in the anterior end of the cell rotating in one direction, while the posterior end rotates in the opposite direction [7]. The physiological significance of this finding is still a matter of debate and a main subject of study.

*T. brucei* is well adapted to *in vitro* culturing, with the procyclic form (PCF) and bloodstream form (BSF) being easily maintained for prolonged periods of time [8–10]. Despite their straightforward culture, imaging free-moving trypanosomes at high resolution has proven to be extremely challenging, given their wide range of motion along the XY and Z axes, and their high swimming speed. Being able to monitor cell movement and cell cycle progression over one or multiple generations would allow us to address various fundamental questions that remain unanswered in the field. Moreover, imaging at high spatial and temporal resolution could allow us to go beyond single species, and address the biology behind interactions between different *Trypanosoma* species.

Current conventional methods to do longitudinal imaging in single parasites *ex vivo* is either using poly-L-lysine-treated coverslips, which result in the parasite’s attachment (partial or total), or the use of gel matrices, which also result in the parasite’s partial or total immobilization. These tools are the gold standard for many studies in the field of *Trypanosoma spp.,* including for example the observation of intraflagellar transport trains, whereby cell immobilization was essential to reveal train movement [11]. Despite their immense value, these methods are incompatible with free-swimming, and the question remains, whether immobilization causes artefacts in the reported observations. Microfluidic devices developed for the study of parasites have been described in several articles. Main methods have been reviewed by Muthinja et al ([12]), and include elastic substrates which allow for the measurement of force transmission during cell migration [13]; nano- and micro-patterns, which can serve as binding sites for motile parasites [14]; micro-pillar arrays that allowing for parasite navigation studies [13]; micro-channels, which have aimed at confining parasites for longitudinal visualization [15]; and organs-on-chip, which have aimed at the replication of *in vivo* parameters for the study of host-parasite interactions (reviewed in [16][17]). A recent device incorporated encapsulation and cultivation of single parasites in emulsion droplets to investigate parasite growth patterns and parasite population heterogeneity [18,19]. While each of these devices comes with specific strengths, with our work we address one common limitation to all: the possibility to maintain free-swimming parasites (single and collective) in the field of view and image them for long periods of time.

In this manuscript, we present a microfluidic approach that aims at preventing *T. brucei* escape from the field of view, using trap geometry to achieve efficient confinement. The produced devices that we called Tryp-Chip, allow the parasites to remain free-swimming, while keeping them within a restricted space for longitudinal imaging for up to 8 hours. Our design allows us to investigate parasite motility and distribution within the device; collective behaviour within the chip and individual traps; and the effect of altering parasite motility by inducible RNAi silencing of paraflagellar rod protein 2 (PFR2), a key component of the paraflagellar rod - a unique structure in the flagellum of kinetoplastid protozoa [20–22]. We envisage that this device could be a valuable tool for high-throughput cell-based assays for *T. brucei* phenotypic characterization/mutant screening, anti-parasitic drug-screening, and the study of individual and collective motility. We see this as a tool for high-temporal and high-spatial resolution imaging, complementary to other valuable methods existing in the field of parasitology.

## Materials and Methods

### Trypanosome cultures and transfection

*Trypanosoma brucei* 2913 procyclic form trypanosomes were routinely grown at 27°C in SDM-79 medium [8] supplemented with 10% (v/v) heat-inactivated foetal calf serum (FCS), 8 mM glycerol (SDMG) and haemin (7.5 mg/ml). For generating the *PFR2^RNAi^* cell line, the pZJMPFR2 plasmid [23] was transfected by Nucleofector technology (Lonza, Italy) [24] in the 2913 cell line expressing the T7 RNA polymerase and the tetracycline repressor [25]. Transfectants were grown in medium with the appropriate antibiotic concentration, and clonal populations were obtained by limiting dilution. For this work, *PFR2^RNAi^* cells were grown in tetracycline for 30 hours to express double-stranded RNA.

### Microfluidic device design and microfabrication

Microfluidic chips were produced by means of standard photo- and soft-lithography (**Figure 1**). Briefly single layer chips were designed with the Clewin™ 5.4 software. The corresponding plastic high resolution photolithography masks were ordered from Micro Lithography Services LTD (Chelmsford, UK). Master molds of heights ranging from 4 to 8 µm were created by means of photolithography on an MJB4 mask aligner using SU8-2005 resin. Height was assessed with a contact profiler (DektakXT, Bruker). After overnight silanization (Trichloro(1H,1H,2H,2H-perfluorooctyl) silane, Sigma-Aldrich) replication of the devices was performed with poly-dimethyl-siloxane (PDMS, Silgard 184; Dow Chemical). Chip bonding was performed using #1.5 coverslips of 22 x 50 mm (Ref 12383138, Fisher Scientific) that were first cleaned by sonication for 20 minutes in isopropanol (Ref 10085103, Fisher Scientific), followed by 20 minutes in Milli-Q® water. The coverslips were then dried with an air gun and stored in sterile conditions until use. Upon obtaining a PDMS slab with an average of 24 chips, we cut sets of 3 chips to assemble on each coverslip. The sets of 3 chips were cut with an octagonal shape, with the corners being cut diagonally. Inlet and outlet holes were generated using a skin hole-puncher with a 1mm diameter. Finally, prepared PDMS slabs and clean coverslips were bonded using an O2 plasma (Cute, Femoto Science, Kr) treatment followed by an incubation at >80°C for 5 minutes. Here the timing was critical as the plasma-induced labile hydrophilicity of the chip greatly helped with the chip loading. There distilled water was injected on each chip via the inlets and stored until use. Chips were used for *T. brucei* culture and imaging on the same day they were produced.

**Figure 1.**
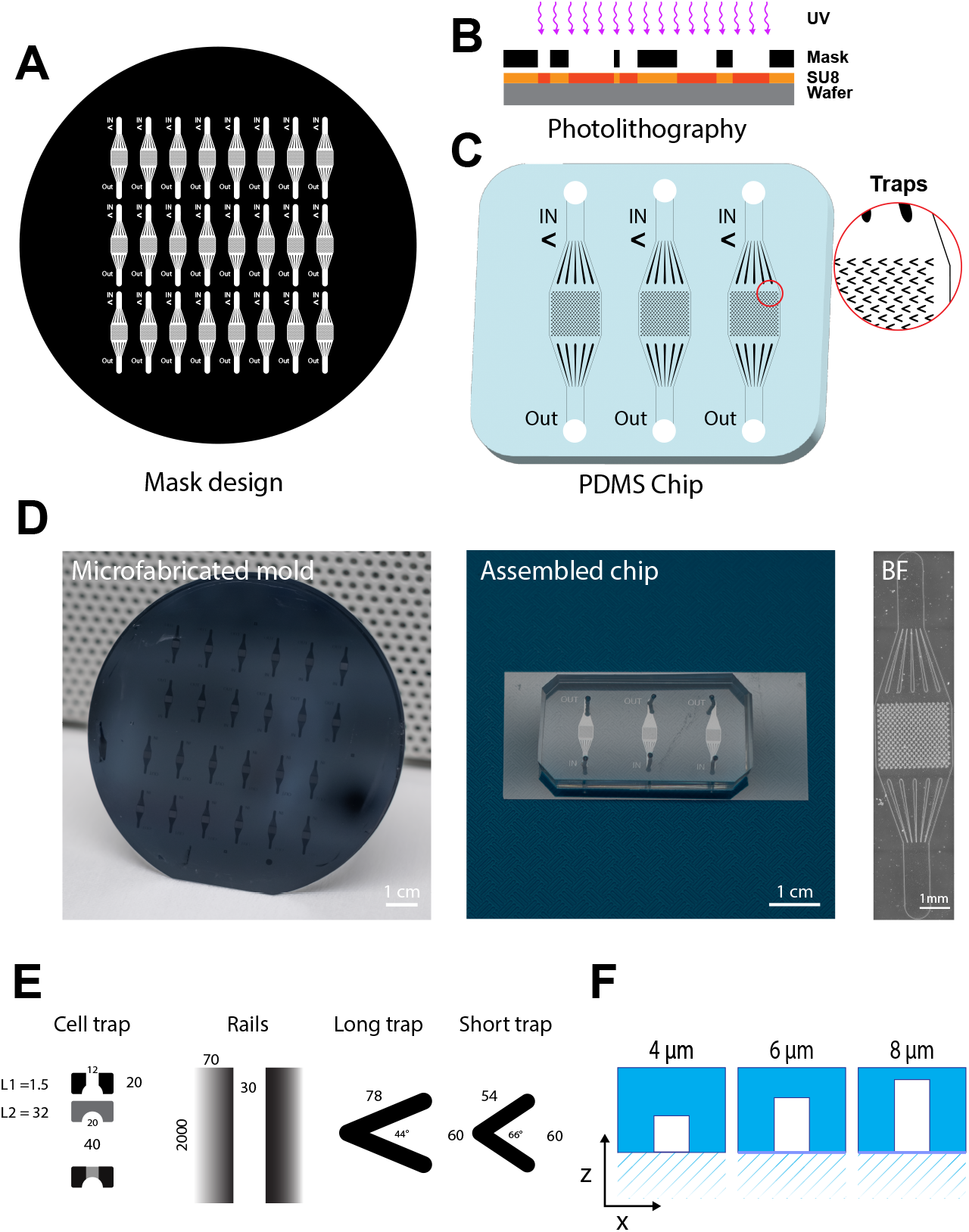
Microfabrication process of Tryp-Chip using photo- and soft-lithography. **A.** A master mold with between 24 and 32 chips was designed using the Clewin^TM^ software, and photolithography on an MJB4 mask aligner. **B.** SU8-2005 photolithography was used for chip microfabrication. This type of lithography uses SU8, a photosensitive negative epoxy (photoresist), which is used to create micro-patterns. It was then exposed to UV light through the mask, to generate the required patterns. The final stages involve hard-baking the photo-resist and curing it. **C-D.** Replication of the devices was performed with poly-dimethyl-siloxane (PDMS), which was poured over the microfabricated mold, and treated under vacuum for at least 30 minutes to remove bubbles, and incubated at 60°C until the PDMS hardened (for a minimum of 4 hours). Once the PDMS mold was ready, chips were cut in sets of 3 (middle panel) for assembly on coverslips. Slabs were cut to achieve an octagonal shape, and inlet and outlet holes were generated using a skin hole-puncher with a 1mm diameter. PDMS slabs and clean coverslips were bonded using O_2_ plasma, and incubated at >80°C for 5 minutes. The individual chips were verified for general accuracy by transmitted light microscopy (right panel). **E.** Each chip contained between 300 and 600 traps of various geometrical shapes and sizes. The 4 geometric designs discussed in this work are hydrodynamic traps, rails, long traps, and short traps. The hydrodynamic cell trap format consists of 2 levels, one with two separate columns forming an open arch (1.5 µm length x 20 µm width), and a second continuous level (32 µm length x 20 µm width). A 12 µm gap on the first level enables the parasites to enter the trap. The rails trap designed was based on 2000 µm (height) x 70 µm (width) columns separated by 30 µm-long tunnels. The long trap design consisted of V-shaped traps with two 78 µm-long arms at a 44° angle. The short trap design consisted of V-shaped traps with 54 µm-long arms at a 66° angle. **F.** For both V-shaped traps, we explored 3 possible arm-heights, namely 4 µm, 6 µm and 8 µm.

### Staining for visualization of *T. brucei* organelles

In order to visualize sub-cellular structures, as proof of concept we stained live *T. brucei* cells with MitoTracker ® Green FM (9074-P, cell signaling technology) to visualize the mitochondrion. A concentration of 200 nM was used in growth medium, and cells were incubated for 30 minutes at 25°C prior to imaging. Hoechst H33342 (62249, ThermoScientific) was used for live imaging of parasite nuclei and kinetoplasts, at a concentration of 1 µg/ml immediately prior to imaging. If Hoechst was added, cells were washed in 1x PBS 3 times for 5 minutes each time, and then resuspended in medium.

### Electron Microscopy

For transmission electron microscopy, cells were prepared as per published protocol [26]. Briefly, cells were fixed in culture medium overnight at 4°C in 2.5% glutaraldehyde – 0.1M cacodylate buffer (pH 7.2). Cells were then washed 3 times in 0.1M cacodylate buffer, and then incubated in 1% tannic acid (in 0.1 M cacodylate buffer) for 30 minutes. Cells were then washed, and post-fixed in 1% OsO4 (in 0.2 M cacodylate buffer) for 1 hour at 25°C. This was followed by 3 washes in distilled water, and post-fixation in 2% OsO4 in water for 1 hour at 25°C. Samples were then washed in distilled water, and then incubated in 1% uranyl acetate (in 25% ethanol) for 30 minutes at 25°C. After serial dehydration in 25%, 50%, 75%, 95% (1x for 10 minutes each) and 100% (3x for 10 minutes each), samples were embedded in Epon resin (propylene oxide 50% + Epon A+B+DMP30 50%) and incubated for a minimum of 48 hours at 60°C for resin polymerization. Ultrathin sections were collected on Formvar-carbon-coated copper grids using a Leica EM ultramicrotome and stained with uranyl acetate and lead citrate. Samples were then visualized on a FEI Tecnai T12 120 kV electron microscope, and images processed using Fiji (ImageJ). Imaged cells include 2913, *PFR2^RNAi^* uninduced, and *PFR2^RNAi^* induced (36h).

### Light Microscopy

Live *T. brucei* parasites were inserted in chips via the inlets immediately prior to imaging. The concentration of parasites ranged from 10^3^ to 10^6^ parasites. An UltraView VOX spinning disc microscope equipped with a Yokogawa CSU-X1 spinning disk head, and two Hamamatsu EM-CCD cameras, was used in bright field mode. For a full view of parasite behaviour in the traps or at their close proximity, we used a 63x oil immersion, 1.4 NA objective. For visualization of the fluorescent signals produced by markers of mitochondria, Golgi apparatus, and endoplasmic reticulum, we used a 100x oil immersion, 1.4 NA objective. For visualization of the full chip, we used a 25x oil 0.8 NA objective. For fast image acquisition, we acquired videos at a maximum speed of 10 fps. The duration of each experiment is specified in the results section. For longitudinal image acquisition spanning minutes, 1 hour or 1-8 hours, short videos spanning 20 frames were acquired once every minute, every 5 minutes or every 15 minutes, respectively. The software used for acquisition was Volocity, and videos were analysed using Fiji [27].

### Statistical analysis

All experiments were repeated in a minimum of triplicates across 5 sets (with each set containing 3 chips). In each chip, a minimum of 200 traps were measured. For full chip colonization measurements, all traps were measured. For experiments where the output refers to individual parasites, a minimum of 1000 parasites was taken into account. For individual parasite tracking and mean square displacement, a minimum of 150 cells per condition were tracked. Statistical analyses were performed in Prism 9 (GraphPad software). Unless otherwise stated, t-tests or one-way ANOVA tests with inter-group comparisons by Tukey *ad hoc* post-tests were performed, and significant results are indicated by * (p < 0.05).

## Results

### Design and generation of various chip models for *T. brucei* constriction analysis

Confining motile unicellular organisms such as *T. brucei* in microenvironments compatible with long term live imaging without inducing exogenous stress calls for the development of dedicated microfabricated device where both geometry and guided fluid flow ensure a proper balance between trapping and survival of the parasite. In order to explore the parameter space of a microfluidic chip that can deliver these feats, we opted for the production of a simple but modular design with a single inlet/outlet and where the geometry of an array of traps can be varied (**Figure 1A-D**). Here we used the standard photolithography process (**Figure 1B**) that allowed for rapid production of various trap designs. The produced chips came with different trap geometries (**Figure 1E**) and trap dimensions (**Figure 1F**). Varying the geometry was meant to explore different trapping concepts including hydrodynamic trapping (cell trapping), friction (rails) and sedimentation (V-shape). Varying the height of the chip was tried out to find the best balance between ease of loading and the maintain in focus of the trapped parasites.

### V-shaped traps result in the highest *T. brucei* trapping efficiency

Previous work has shown that *T. brucei* parasites display different navigational skills that depend on the geometrical pattern and separation of micropillars (or obstacles) present in their environment (reviewed in [12]). Here, our aim was to physically constrain *T. brucei* procyclic parasites for long-term imaging, while allowing them to remain capable of moving the flagellum freely (as opposed to immobilization strategies that rely on agar or chemical adhesion via poly-L-lysine or other substrates). The average length of a procyclic *T. brucei* is between 20 µm (uniflagellated cells) and 25 µm (cells about to divide, with two flagella) [28] and the average width was 2.5 µm. We began by exploring various traps with different geometrical properties to determine the one resulting in the highest trapping efficiency to spatially constrict *T. brucei* motility for long-term imaging. The first analysis we performed consisted in determining the percentage of traps of the 4 shapes described in **Figure 1E** that resulted in retention of one or more parasites for a minimum of 30 seconds (**Figure 2A**). Then, we went on to determine *T. brucei* behaviour within the trap that contributed (or not) to retention, by measuring the total retention time up to a maximum of 30 seconds (**Figure 2B**). We defined this time threshold based on our observations on parasite distribution time, to differentiate passive/random entry into traps, and active permanence (i.e. parasites entering and remaining in the trap, as opposed to randomly entering and exiting). Retention was measured in a minimum of 5 chip sets (containing 3 chips each: 15 chips in total) distributed across 3 separate experiments, and a minimum of 100 traps per chip. For this initial investigation, the height (z-axis) we tested for the rails and the V-shaped traps was of 8 µm.

**Figure 2.**
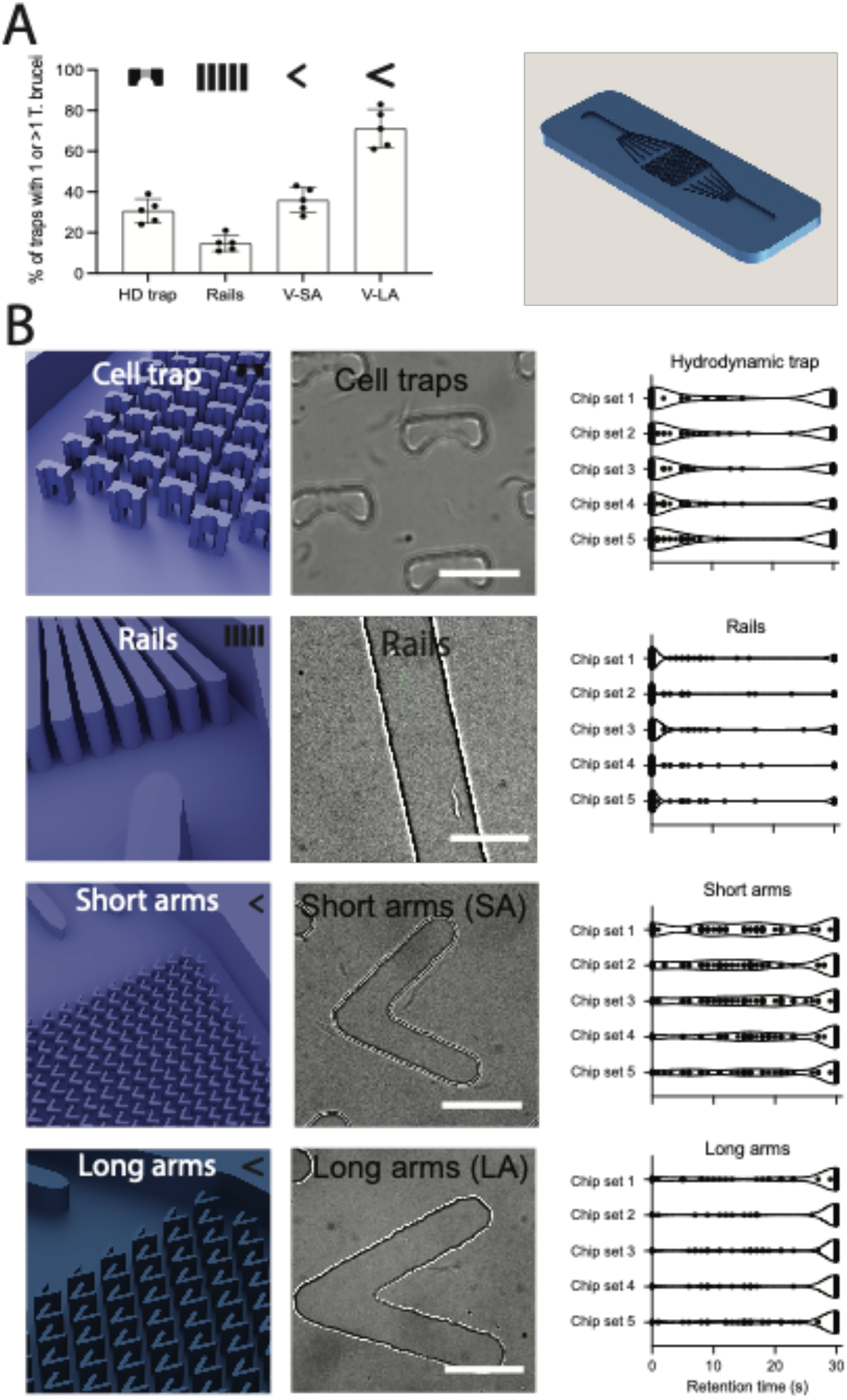
Trapping efficiency of different geometrical trap designs. Each design was tested for *T. brucei* trapping efficiency and retention time. **A.** We began by quantifying the percentage of traps that resulted in parasite retention for a minimum of 30 seconds. The hydrodynamic trap showed between 24 and 39% retention across 5 repeats (average: 30.6%, SD: 5.9) across 1500 traps measured in total. The rails showed between 11 and 21% retention (average: 15%, SD: 4.1) across 300 rails measured in total. The V-shaped traps with short arms showed between 28 and 43% retention (average: 36%, SD: 6.2) across 4500 traps measured in total, and the V-shaped traps with long arms showed between 61 and 83% retention (average: 71.2%, SD: 9.4) across 4500 traps measured in total. The right panel shows the overall design used for all chips, with only the traps in the central squared region changing from one design to another. **B.** The left-most panels show a 3D view of the traps in each design. The middle panels show bright field images of the traps, acquired with 100x magnification, populated by *T. brucei.* Scale bar is 25 µm. The right-most panels show truncated violin plots showing highly-resolved parasite behaviours within traps. For clarity, results for one chip are shown as an example. In the hydrodynamic traps, among all invaded traps, between 37 and 59% were not invaded at any point. Between 8 and 29% were invaded and exited within the first 10 seconds, between 1 and 8% were invaded and exited within the next 10 seconds (20 seconds total) and only between 24 and 41% were invaded and remained invaded by 30 seconds. In the rail design, between 61 and 76% of rails did not result in trapping at any point. Within the first 10 seconds, between 8 and 16% of rails showed parasite retention. In the next 10 seconds (i.e. 20 seconds total) between 1 and 3% of rails led to temporal trapping, and only between 10 and 29% continued to show trapping after 30 seconds. In the short-arm V-shaped trap, between 6 and 19% of traps were unsuccessful in trapping parasites at any time. Then, between 12 and 25% of traps were invaded and exited within the first 10 seconds. Between 23 and 38% of traps remained invaded after 20 seconds. Between 23 and 49% were invaded, and continued to be so within 30 seconds. Finally, between 5 and 13% of long-arm V-shaped traps were not invaded at any time point. Between 2 and 15% were invaded and exited within the first 10 seconds of imaging; between 7 and 19% remained invaded within 20 seconds of imaging, but parasites entered and exited the trap within this time. Between 63 and 83% of traps were invaded and remained so after 30 seconds of imaging. V-shaped traps with long arms were, overall the most efficient for parasite trapping and were selected for the rest of the work presented here. All experiments are the result of measurements across 1500 traps or 300 rails in 15 chips. All data for Figure 2 are found in **Supp. Table 1**.

First, we tested the hydrodynamic cell trap (consisting of 2 levels, one with two separate columns forming an open arch (1.5 µm length x 20 µm width), and a second continuous level (32 µm length x 20 µm width) – a 12 µm gap on the first level enables the parasites to enter the trap. Hypothetically, the geometrical form of the trap would make parasite escape complicated. We observed that between 24 and 39% of traps were successfully invaded (mean: 30.6, SD: 5.94) (**Figure 2A**). High-temporally resolved (i.e. image acquisition rate) investigation showed that most traps were not invaded (up to 59%), or led to escape within the first 10 seconds after invasion (up to an additional 29%) (**Figure 2B, top row**). Our main explanation for this low ‘retention’ was low statistical likelihood of parasite entry given the small surface of the gap compared to the rest of the environment in the trap. It was less likely for parasites to find the 12 µm gap to enter the trap, despite showing exploratory behaviour in the trap’s surroundings (**Video 1**).

Next, we tested parasite retention in rails (2000 µm length x 70 µm width with a 30 µm gap between rails). In the figure, the rail is shown from the top. We hypothesized that *T. brucei* could be trapped by increasing friction along its length. Namely, that parallel rails matching the average *T. brucei* length (25 µm) would succeed in trapping parasites by generating friction along the parasite’s long axis. Unfortunately, this relative positioning was rare, and in a parallel position relative to the rail, up to 76% of rails resulted in no trapping. On average, less than 16% of rails result in *T. brucei* trapping for more than 30 seconds (**Figure 2A**). Further investigation showed that although almost all parasites enter the rails, the vast majority do not stop along the rail. Rather, they use the rail as a track to navigate quickly in a position parallel to the rail (**Figure 2B, second row** and **Video 2**).

We then hypothesized that a V-shaped trap could succeed in constricting trypanosome motion both in the x- and y-axis of any parasite successfully entering the trap. The idea was to offer areas of low flow orthogonal to the inlet/outlet flow axis. We first tested V-shaped traps, which we call ‘short arms (SA)’, with 54 µm-long arms and 60° aperture. Opposite to our observations in the hemodynamic cell trap and the rails, parasites here successfully entered the trap in many cases (in all chips tested and all replicates). However, after showing exploratory behaviour within the trap, many parasites escaped within 10-20 seconds (**Figure 2B, third row** and **Video 3-4**). Nevertheless, this design resulted in 28-43% of traps (mean: 36%, SD: 6.2) successfully retaining *T. brucei* parasites for at least 30 seconds (**Figure 2A**).

We next considered that increasing the length of the arms and closing the angle of aperture could increase parasite retention in the trap. We designed a second V-shape with 78 µm arm length and 44° aperture, which we called ‘long-arms (LA)’. This design resulted in a 40% increase in parasite retention relative to the SA trap (average: 71.2%, SD: 9.4) (**Figure 2A**). In the V-shaped LA design, most parasites entered the trap, and remained within it for 30 seconds (up to several hours) (**Figure 2B, bottom row**), (**Video 5 and 6**).

### Trap height influences *T. brucei* trapping efficiency

Having defined the most efficient geometrical shape for *T. brucei* trapping, we observed in the original 8 µm height design, that while parasites were spatially confined in the x- and y-axes of the V-LA trap, their motility along the z-axis remained large enough to still make imaging challenging. The cell body and the flagellum were constantly coming in and out of focus, impeding high-resolution observations. Moreover, we observed that most traps retained multiple parasites, rather than single parasites in multiple traps. While this was not a negative outcome, our target was to find the height and parasite concentration that was best suited for single cell trapping and single cell imaging.

We therefore compared parasite trapping properties in chips of 4 µm, 6 µm and 8 µm of height (**Figure 3A**). These values were chosen based on the *T. brucei* average width which is between 2 and 4 µm. according to the stage of the cell cycle. With the aim of achieving single cell trapping, we also tried various parasite concentrations (10^3^, 10^4^, 10^5^ and 10^6^ parasites /µl) (**Figure 3B**). In devices with a 4 µm height (**Figure 3B, left panel**), the dominant observation was that parasites were difficult to insert into the chip. Upon insertion, the majority of traps (86.5%) were either empty (at a concentration of 10^3^ parasites/µl), or had mostly lysed parasites (at concentrations of 10^4^-10^6^ parasites/µl) (77.7, 75.1 and 87.3% respectively). We concluded that a height of 4 µm was deleterious for parasite survival.

**Figure 3.**
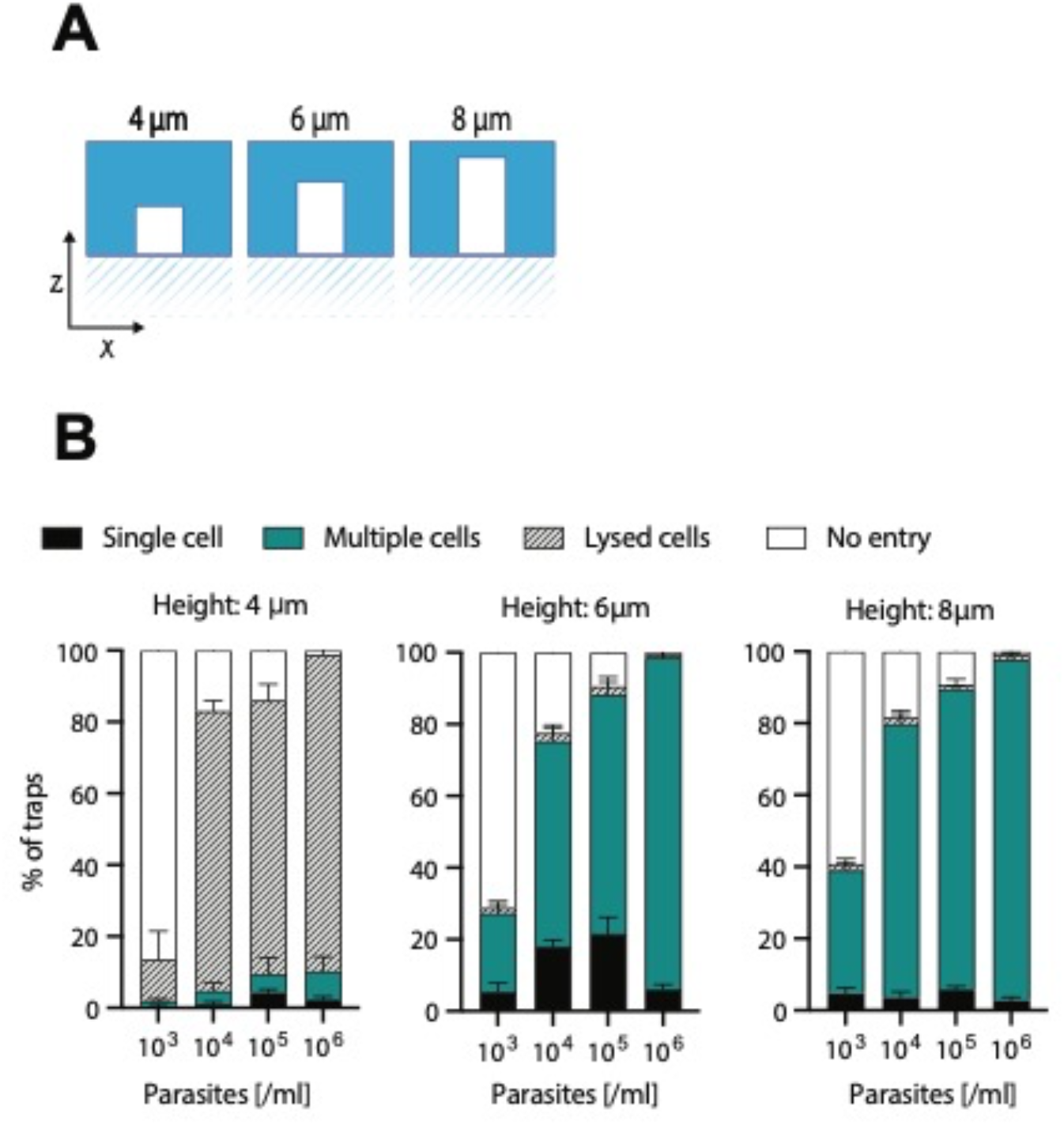
Trapping efficiency in V-shaped traps according to arm height. V-shaped traps with long arms were designed with 3 different arm height to test whether this factor was important for trapping efficiency. **A.** Arm heights tested were 4 µm, 6 µm and 8 µm. **B.** Four parasite concentrations were tested for each of the 3 arm heights, namely, 10^3^, 10^4^, 10^5^, and 10^6^ parasites/ml. **Left panel: 4 µm height.** At a concentration of 10^3^ parasites/ml, an average of 86.5% (SD 5.95) of traps were not invaded, while an average of 11.7% (SD 6.1) showed lysed cells. Only the remaining 2% of traps were invaded by either single parasites (0.7%, SD 0.72) or multiple parasites (1.2%, SD 0.77) per trap. At a higher concentration, namely 10^4^ parasites/ml, an average of 17% (SD 4.4) of traps were not invaded, while 77.7% (SD 3.6) showed lysed parasites. Only 5.3% of traps were invaded either by single parasites (1.1%, SD 0.83) or multiple parasites (4.2%, SD 2.62). At a concentration of 10^5^ parasites/ml, a similar result was obtained. Namely, an average of 14% (SD 6.7) of traps were not invaded, while 75.1% (SD 4.6) showed lysed parasites. Only 10.9% of traps were invaded either by single parasites (5.2%, SD 2.4) or multiple parasites (5.7%, SD 3.7) per trap. A similar situation was observed at a concentration of 10^6^ parasites/ml, namely, 2.1% (SD 1.4) of traps were not invaded, while 87.3% (SD 4.8) showed lysed cells. 10.5% of traps were successfully invaded either by single (2.3%, SD 1.5) or multiple (8.2%, SD 4.3) parasites per trap. Altogether, at all concentrations, the dominant observation was that in traps with a 4µm height, was that parasite lysis heavily occurs. **Middle panel: 6 µm height.** At a concentration of 10^3^ parasites/ml, an average of 72.5% (SD 4.5) of traps were not invaded, while an average of 1.1% (SD 0.8) showed lysed cells. Conversely, 26.3% of traps were invaded either by single parasites (5.8%, SD 2.8) or multiple parasites (20.5%, SD 3.5) per trap. At a concentration of 10^4^ parasites/ml, an average of 22.9% (SD 4.9) of traps were not invaded, while 1.8% (SD 1.0) showed lysed parasites. Conversely, 75.3% of traps were invaded either by single parasites (18%, SD 2.1) or multiple parasites (57.3%, SD 4.9). At a concentration of 10^5^ parasites/ml, an average of 9.5% (SD 4.1) of traps were not invaded, while 2.4% (SD 1.2) showed lysed parasites. Conversely, 88.1% of traps were invaded either by single parasites (21.9%, SD 3.5) or multiple parasites (66.3%, SD 2.9) per trap. Finally, at a concentration of 10^6^ parasites/ml, 0.7% (SD 0.8) of traps were not invaded, while 0.7% (SD 0.6) showed lysed cells. Conversely, 98.5% of traps were successfully invaded either by single (6.1%, SD 0.96) or multiple (92.5%, SD 0.92) parasites per trap. Altogether, the increase in height by 2µm overcame the challenge of cell lysis posed by the 4µm-height traps. **Right panel: 8 µm height.** At a concentration of 10^3^ parasites/ml, an average of 59.9% (SD 4.3) of traps were not invaded, while an average of 1.3% (SD 1.1) showed lysed cells. Conversely, 38.7% of traps were invaded either by single parasites (5.1%, SD 1.3) or multiple parasites (33.6%, SD 3.4) per trap. At a concentration of 10^4^ parasites/ml, an average of 18.8% (SD 4.0) of traps were not invaded, while 1.5% (SD 0.5) showed lysed parasites. Conversely, 79.7% of traps were invaded either by single parasites (3.5%, SD 1.4) or multiple parasites (76.1%, SD 3.6). At a concentration of 10^5^ parasites/ml, an average of 9.9% (SD 3.3) of traps were not invaded, while 0.8% (SD 0.9) showed lysed parasites. Conversely, 89.3% of traps were invaded either by single parasites (5.8%, SD 1.5) or multiple parasites (83.5%, SD 2.9) per trap. Finally, at a concentration of 10^6^ parasites/ml, 1.3% (SD 0.8) of traps were not invaded, while 1.5% (SD 0.9) showed lysed cells. Conversely, 97.3% of traps were successfully invaded either by single (2.7%, SD 1.6) or multiple (94.5%, SD 1.7) parasites per trap. All experiments are the result of measurements in 15 chips, and 1500 traps.

In devices with a 6 µm height (**Figure 3B, middle panel**), we observed that at a concentration of 10^3^ parasites/µl, the majority of traps (72.5%) were empty. By contrast, using higher parasite concentrations (10^4^-10^6^ parasites/µl) resulted in most of the traps being populated by multiple (i.e. >3) parasites (57.3, 66.3, and 92.5% respectively). Moreover, parasite lysis was greatly diminished compared to the one observed in 4 µm-height chips (less than 2%). When examining trypanosome distribution in traps, 6 µm-high chips inoculated with concentrations of 10^4^-10^5^ parasites/µl showed the highest rates of single cell retention (18 and 21.9% respectively) observed amongst all trap configurations, trap heights, and parasite concentrations tested so far.

With devices of an 8 µm height (**Figure 3B, right panel**), we observed that at a concentration of 10^3^ parasites/µl, the majority of traps (59.9%) were empty, similar to what we observed in the two other devices. At higher parasite concentrations (10^4^-10^6^ parasites/µl), most traps were populated by multiple parasites (76.1, 83.5, and 94.5% respectively). Importantly, parasite lysis was minimal (less than 1.8%). However, the rates of single cell trapping at 10^4^-10^6^ parasites/µl were very low (5.8% at best).

Altogether, we determined that although the general parasite retention efficiency was similar in V-shapes LA traps of 6 µm and 8 µm of height, more individual/single parasites in 6 µm-high traps remained in focus throughout long periods of time. Moreover, we defined that 10^4^ and 10^5^ parasites/ µl were the most suitable concentrations for parasite analysis in our microfluidic devices. At these concentrations, 6 µm-high traps were the most successful in single parasite retention. For the remaining work presented here, we generated all devices with 6 µm of height, and used concentrations of 10^4^ and 10^5^ parasites/ µl (unless otherwise indicated).

### Trap density influences *T. brucei* trapping efficiency

Having defined trap geometry and parasite density suitable for in-focus imaging, we then compared the impact of trap density on trapping efficiency (**Figure 4**), using high (i.e. ∼600 traps in a 4mm^2^ area, distributed in 22 columns x 28 rows) (**Figure 4A**) or low density of traps (i.e. ∼300 traps in a 4mm^2^ area, distributed in 19 columns x 16 rows) (**Figure 4E**). To investigate the effect of trap density on trapping efficiency, we divided the total trap region into 12 quadrants, separated into 3 columns (1-3) and 4 rows (A-D) with equal surface and the same numbers of traps (as shown in **Figures 4B and 4F**). In the schematic, row A is the one closest from the inlet where parasites are inserted, while row D is the one furthest away (and closest to the outlet). Upon measuring the percentage of traps which successfully captured *T. brucei*, we noticed that in chips with a high density of traps, there was a consistent gradient in invasion with decreasing numbers of traps being invaded as the distance from the inlet increased. This was reproduced in 15 separate chips. More precisely, row A had an average of 90.6% of traps invaded, row B an average of 88.3% of traps invaded, row C an average of 52.1% of traps invaded, and row D an average of only 19.2% of traps invaded. Notably, the invasion by column was not significantly different. We hypothesize this is largely due to the fact that the design includes channels that evenly distribute the fluid inserted into the inlet, before it reaches the chip region that harbours the traps.

**Figure 4.**
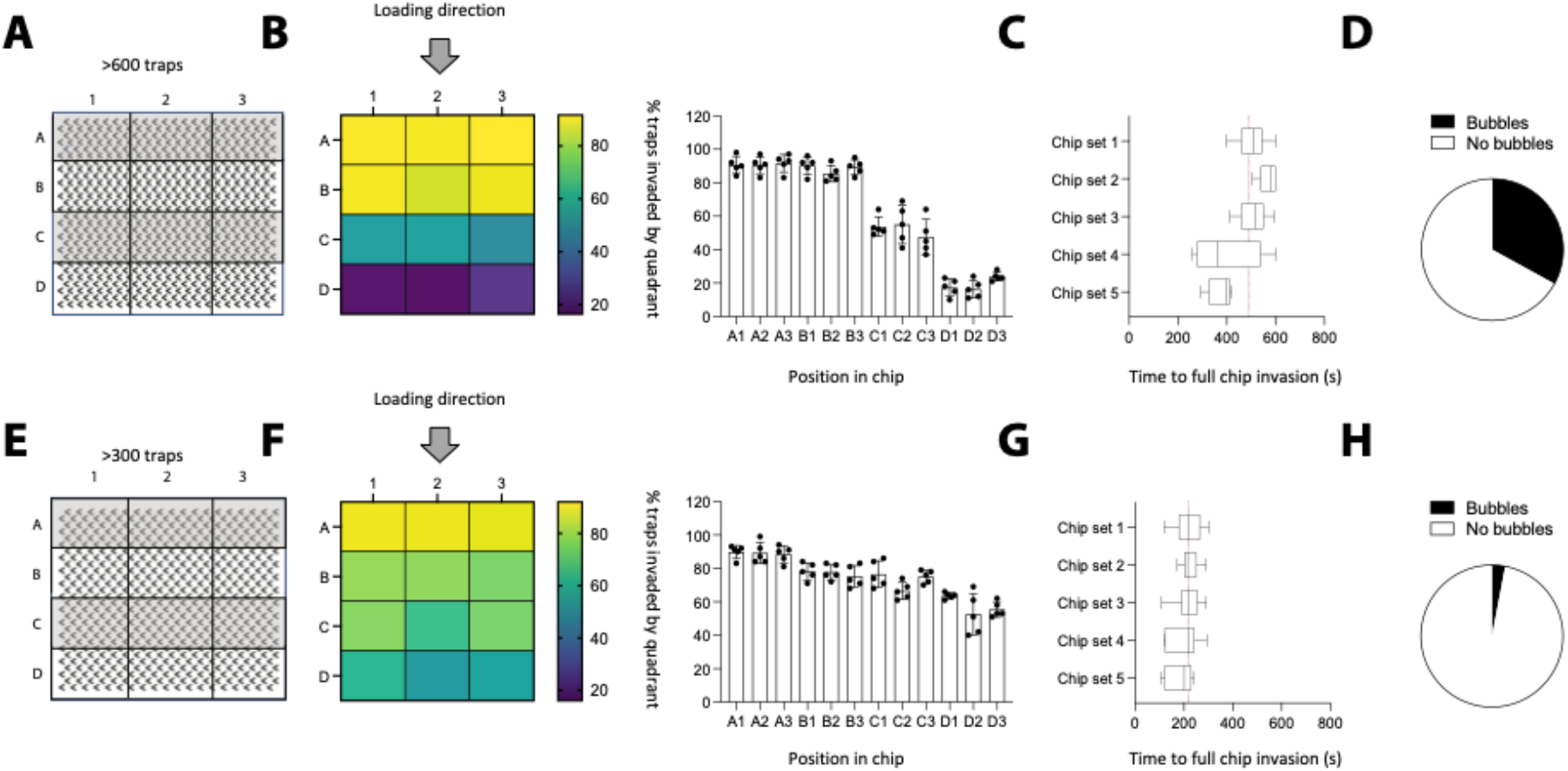
Trap density impacts trapping efficiency of Tryp-Chip. We compared the trapping efficiency of chips with relatively high (i.e. 600 traps per chip, or 150 traps/mm^2^) or low (i.e. 300 traps per chip, or 75 traps/mm^2^) trap densities. **A-B.** For high density chips, we divided the total chip region into 12 quadrants, separated into 3 columns (1-3) and 4 rows (A-D) with equal surface and the same numbers of traps per quadrant. The inlet allowing parasite influx was located above the A row. A series of channels designed between the inlet and the traps were used in our design to ensure uniform distribution of cells upon their arrival to the chip region following insertion. **B. Left panel.** Heat map represents the average percentage of invaded traps per quadrant. **Right panel.** Histograms show the individual results and SDs of 5 chip sets (i.e. 15 chips), whereby a total of 1500 traps were imaged per set. For the high density chips, the following results show the average percentage of invaded traps: in the upper-most row, closest to the inlet, quadrant A1: 90.6%, A2: 90.4% and A3: 91.6%. In the second row: B1: 90.2%, B2: 85.4% and B3: 89.2%. In the third row: C1: 53.8%, C2: 55% and C3: 47.6%. In the bottom-most row, D1: 17.4%, D2: 16.4%, and D3: 23.8%. This represents a 71% decrease in invaded traps between the top-most row, and the bottom-most row. This difference is statistically significant (p < 0.001). **C.** In addition to percentage of invaded traps, we also calculated the time taken by >5 parasites to reach row D. In the multiple chip sets we evaluated, we found that the median time to full invasion was 520.5 s (SD: 65.5), 572 s (SD: 30.1), 518.5 s (SD: 59.2), 398 s (SD: 131.7) and 399.5 (SD: 47.2). **D.** We assessed whether high trap density influenced fluid distribution within the chip by assessing the percentage of traps with bubbles – a significant hindrance for parasite evaluation and trapping efficiency. Altogether, 33% of chips had bubbles. **E-H.** Next, we evaluated low-trap-density chips. **F. Left panel.** Heat map represents the average percentage of invaded traps per quadrant. **Right panel.** Histograms show the individual results and SDs of 5 separate experiments (i.e. 15 chips). For the low density chips, the following results show the average percentage of invaded traps: in the upper-most row, closest to the inlet, quadrant A1: 89.8%, A2: 89.2% and A3: 88.4%. In the second row: B1: 78.2%, B2: 77.8% and B3: 75.2%. In the third row: C1: 76.6%, C2: 66.6% and C3: 75%. In the bottom-most row, D1: 63.5%, D2: 52.6%, and D3: 55.4%. This represents a 30% decrease in invaded traps between the top-most row, and the bottom-most row. This difference is statistically significant (p < 0.05), yet not as pronounced as the one observed in high-density-traps. **G.** In addition to percentage of invaded traps, we also calculated the time taken by >5 parasites to reach row D. In the 5 chip sets we evaluated, we found that the median time to full invasion was 219 s (SD: 48.3), 215.5 s (SD: 33.4), 220.5 s (SD: 52.9), 218 s (SD: 56.9) and 204 (SD: 48.8). **D.** We assessed whether high trap density influenced fluid distribution within the chip by assessing the percentage of traps with bubbles and found that altogether, 3% of chips had bubbles. All data for Figure 4 is found in **Supp. Table 3**.

This gradient turned out to be less pronounced in chips with lower trap densities (**Figure 4F**). Row A (closest to the inlet) had an average of 89.1% of traps invaded, row B an average of 77% of traps invaded, row C an average of 72.7% of traps invaded, and row D an average of 57.2% of traps invaded. While the difference between row A and row D is significant in both types of chips (high and low trap densities), the lower density chip showed more even parasite distribution across the traps.

We also noticed that the time for full chip colonization (i.e. the time for at least 5 parasites to reach row D) was significantly different between the two conditions. In chips with a high trap density, the average colonization time was of 478.5 seconds (SD 100.6) (**Figure 4C**), while in chips with a low trap density, the average colonization time was of 210 seconds (SD 48.5), so more than twice faster (**Figure 4G**). Our live observations suggested on one hand that high trap densities were hindering *T. brucei* navigation. On the other hand, as shown in **Figure 3B**, *T. brucei* tend to preferentially invade traps in multiple numbers (i.e. swarm). We will further explore this in **Figure 8**. Finally, we also observed that in high density traps, the probability of inserting air bubbles is higher (33% of quadrants per chip have bubbles) (**Figure 4D**) than in low density traps (where only 3% of quadrants show bubbles) (**Figure 4H**).

Based on these results, we established that the prototypes we would use would be low density (∼300-400 traps) chips populated with V-shaped traps with 78 µm-long arms, 44° aperture and 6 µm height, with optimal concentrations of 10^4^-10^5^ cells/ µl.

### Microfluidic devices allow the characterization of mutant parasites with altered motility

To determine whether our prototype would allow high-throughput characterization of *T. brucei* behaviour, we used a cell line where motility was drastically reduced following tetracycline-inducible RNAi knockdown of the *PFR2* gene [23]. The *PFR2^RNAi^* mutant has a greatly diminished paraflagellar rod and has been reported to display highly reduced motility [20,22]. For our characterization, we analysed 3 cell lines in parallel, namely, the parental line 2913, the uninduced *PFR2^RNAi^* (-Tet) line, and the 30 hour-induced *PFR2^RNAi^*(+Tet) line. **Figure 5A** shows fluorescence microscopy images using the antibody 2E10 [29], which labels the paraflagellar rod. Here, we observe that the 2913 and the *PFR2*^RNAi^ (-Tet) lines have a fully labeled paraflagellar rod, while the *PFR2*^RNAi^ (+Tet) line has an intermittent labeling, and an accumulation (blob) at the tip of the flagellum. This is consistent with the previously defined phenotype for this line [20,22]. **Figure 5B** shows electron microscopy images where white arrows point at the paraflagellar rod in each line. Here, we observe that the 2913 and the *PFR2*^RNAi^ (-Tet) lines have a relatively unaltered paraflagellar rod, while the *PFR2*^RNAi^ (+Tet) line has a greatly diminished paraflagellar rod.

**Figure 5.**
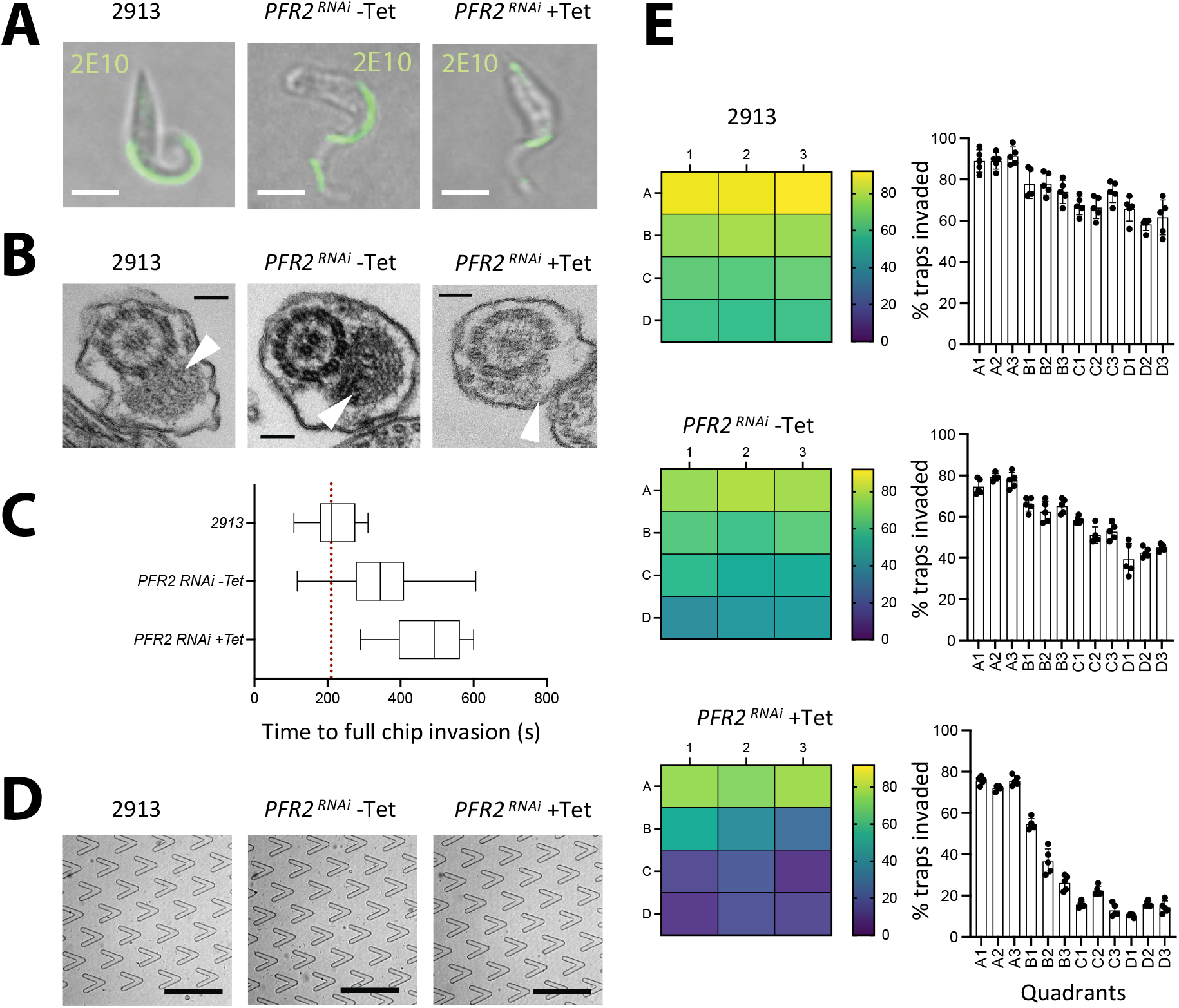
Tryp-Chip allows the characterization of motility mutants. Tryp-Chip was used to characterize the invasion capacity of *PFR2^RNAi^* mutants. **A.** Bright field and fluorescence microscopy images showing paraflagellar rod labeling using antibody 2E10 (scale bar, 10 µm). The 2913 parental line, and the *PFR2*^RNAi^ (-Tet) control line have a fully labeled paraflagellar rod, while the *PFR2*^RNAi^ (+Tet) line has an intermittent labeling, and an accumulation (bleb) at the tip of the flagellum. **B.** Transmission electron microscopy images (scale bar, 100 nm) of the 2913 parental line, and *PFR2*^RNAi^ (-Tet) and *PFR2*^RNAi^ (+Tet) lines. White arrows point at the paraflagellar rod in each line. The 2913 and the *PFR2*^RNAi^ (-Tet) lines have a relatively unaltered paraflagellar rod, while the *PFR2*^RNAi^ (+Tet) line has a greatly diminished paraflagellar rod. **C.** Average time for full chip invasion (i.e. >5 parasites reaching row D as described in Figure 4. The 2913 parental line showed a median invasion time of 210 seconds, while the *PFR2*^RNAi^ (-Tet) control line showed a median invasion time of 341 seconds. The *PFR2*^RNAi^ (+Tet) line showed showed a median invasion time of 496 seconds. In all cases, the difference between all 3 lines are statistically significant (p < 0.01). **D.** Bright field images of parasite presence in row D 600 seconds after insertion into the inlet (scale bar, 250 µm). **E. Top panels.** Refer to Figure 4F panels, previously discussed, and included in this figure for direct comparison with the *PFR2*^RNAi^ induced and non-induced conditions. **Middle panel, left.** Heat map represents the average percentage of invaded traps per quadrant corresponding to the *PFR2*^RNAi^ (-Tet) control parasite line. **Middle panel, right.** Bar chart shows the individual results and SDs of 5 separate experiments (i.e. 15 chips). A1: 74.6%, A2: 79.2% and A3: 77.6%. In the second row: B1: 66%, B2: 62.4% and B3: 65.2%. In the third row: C1: 58.4%, C2: 51.2% and C3: 52.8%. In the bottom-most row, D1: 39.4%, D2: 42.6%, and D3: 45.0%. This represents a 35% decrease in invaded traps between the top-most row, and the bottom-most row. This difference is statistically significant (p < 0.001). **Bottom panel, left.** Heat map represents the average percentage of invaded traps per quadrant corresponding to the *PFR2*^RNAi^ (+Tet) parasite line. **Bottom panel, right.** Bar chart shows the individual results and SDs of 5 separate experiments (i.e. 15 chips). A1: 75.8%, A2: 72.0% and A3: 75.6%. In the second row: B1: 54.6%, B2: 36.6% and B3: 26.2%. In the third row: C1: 15.6%, C2: 22.4% and C3: 12.8%. In the bottom-most row, D1: 10.4%, D2: 16%, and D3: 14.4%. This represents a 60% decrease in invaded traps between the top-most row, and the bottom-most row. This difference is statistically significant (p < 0.001). All data for Figure 5 is found in **Supp. Table 4**.

We began by determining whether the time for full chip invasion (i.e. reaching quadrant D) was equal for all lines. Measurements were done across 50 chips per line (in 3 separate experiments) (**Figure 5C**). This revealed that the time for chip colonization was significantly different, with the 2913 line taking a median of 210 seconds, the *PFR2*^RNAi^ (- Tet) taking a median of 341 seconds, and the *PFR2*^RNAi^ (+Tet) taking a median of 496 seconds. **Figure 5D** shows parasite presence in row D at 600 seconds for all lines. Next, we characterized invasion dynamics by quadrant using the 3 parasite lines, as previously explained in **Figure 4**.

Upon investigation of *T. brucei* distribution, we found that chip invasion by the 3 samples shows significantly different dynamics as determined at 600 seconds following insertion via the inlet. We previously discussed the results obtained in the parental control (**Figure 4F**), and show repeats run in parallel to the *PFR2^RNAi^* for comparison (**Figure 5E top panels**). In the case of *PFR2*^RNAi^ (-Tet), an average of 60% of traps were invaded, with row A showing the highest invasion (77.1%), and row D the lowest (42.3%) (**Figure 5E, middle panel**). Finally, the chips used to investigate the induced *PFR2*^RNAi^ (+Tet) had on average only 36% of traps invaded, with row A showing the highest invasion (74%, similar to the non-induced control), and row D the lowest (13.6%, three times less than the controls) (**Figure 5E, bottom panel**). The *PFR2*^RNAi^ (+Tet) parasites were both, less capable of entering the V-shaped traps across the full chip, and less able to traverse the full chip, resulting in an accumulation of parasites in rows A and B. This emphasizes the relevance of the paraflagellar rod for navigation and sensing which has been previously proposed but so far not tested.

Further investigation of 2913 parental *T. brucei* behaviour in the proximity of traps allowed us to define 4 main behaviours (**Figure 6A**): 1) parasite entry into the V-shaped trap, followed by exit within the first 30 seconds. Parasites then would either continue to surround the trap, or move away from the field of view. In the parental control line, this behaviour was observed in a minority (3.3%) of parasites; 2) parasite attachment to or contact with other trap locations different to the space within the V-shaped trap. This was observed in 18.1% of parasites; 3) unsuccessful interaction, namely, the parasite does not enter the V-shaped trap. This was observed in 19.9% of parasites; and 4) interaction with the corner, whereby parasites enter the V-shaped trap, and preferentially and persistently stay in the proximity of the corner. This behaviour was dominant, with over 58.7% of cells displaying interactions with the trap corner (**Figure 6B**), irrespective of whether the entrance to the trap was with the flagellum leading or otherwise.

**Figure 6.**
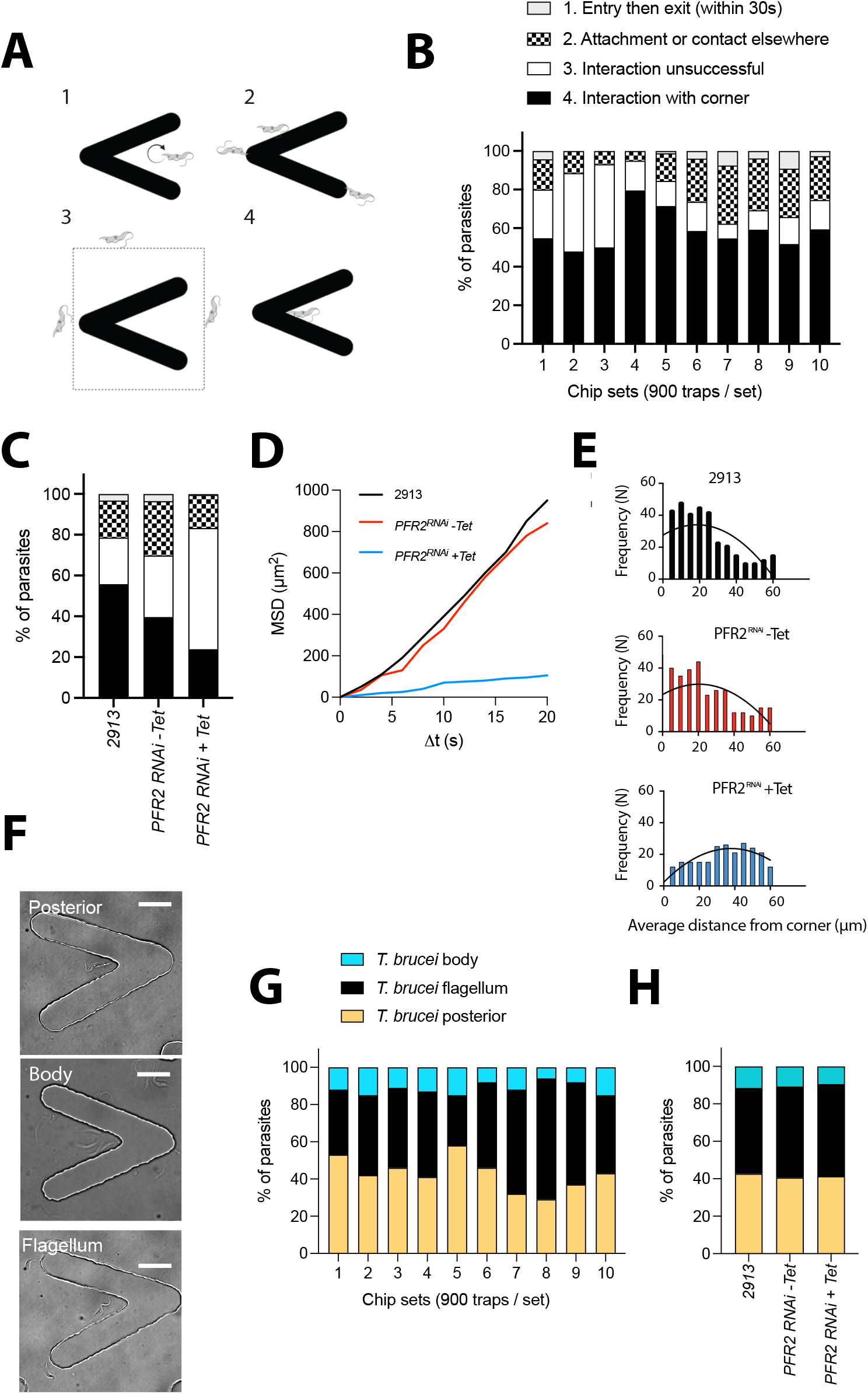
Tryp-Chip allows the characterization of displacement and parasite-trap interactions. **A.** Investigation of *T. brucei* interactions with traps in Tryp-Chip allowed us to define 4 main behaviours: 1. Parasite entry into the V-shaped trap, followed by exit within the first 30 seconds. 2. Parasite attachment to or contact with other trap locations different to the space within the V-shaped trap. 3. Unsuccessful interaction, whereby the parasite does not enter the V-shaped trap, nor does it interact with it. 4. Interaction with the corner, whereby parasites enter the V-shaped trap, and preferentially and persistently stay in the proximity of the corner. **B.** Quantification of the 4 main behaviours in 10 different chip sets demonstrated that in multiple sets of the parental line, on average, 3.3% of parasites showed behaviour 1 (entry then exit); 18.1% showed behaviour 2 (interactions elsewhere); 19.9% showed behaviour 3 (no interaction); and 58.7% showed behaviour 4 (interaction with the corner of the trap). **C.** Quantification of the 4 main behaviours across the parental and *PFR2*^RNAi^ induced and non-induced conditions. 3.5% of *PFR2*^RNAi^ (-Tet) parasites showed behaviour 1 (entry then exit); 26.2% showed behaviour 2 (interactions elsewhere); 30.1% showed behaviour 3 (no interaction); and 40.3% showed behaviour 4 (interaction with the corner of the trap). Conversely, 0.5% of *PFR2*^RNAi^ (+Tet) parasites showed behaviour 1 (entry then exit); 16% showed behaviour 2 (interaction elsewhere); 59% showed behaviour 3 (no interaction); and 24.5% showed behaviour 4 (interaction with the corner of the trap). **D.** Mean squared displacement (MSD) of the parental line 2913 is shown in black; *PFR2*^RNAi^ (-Tet) in red and *PFR2*^RNAi^ (+Tet) in blue. Results are the average of at least 20 parasites tracked per chip, in 3 separate chips. **E.** Frequency of parasite positioning at various distances from the trap corner. The parental 2913 (black bars) and *PFR2*^RNAi^ (-Tet) parasites (red bars) preferentially localize within 0-20 µm (i.e one body length) from the trap corner. *PFR2*^RNAi^ (+Tet) parasites are scattered across the trap, with a higher frequency occurring between 30 and 60 µm from the trap. **F-G.** Parasite positioning and interaction with the trap. **F.** Bright field images representative of the parasite posterior interacting with the trap **(top panel)**, the parasite body interacting with the trap **(middle panel)** or the parasite flagellum interacting with the trap **(bottom panel). G.** Multiple chip sets showing parasite orientation for trap interaction. An average of 42.7% (SD 8.8) parasites were found to interact with the posterior; 45.8% (SD 10.8) with the flagellum, and 11.5% (SD 3.2) with the parasite body. **H.** Compared to the control parental line, an average of 40.3% (SD 3.2) *PFR2*^RNAi^ (-Tet) parasites were found to interact with the posterior; 48.6% (SD 6.4) with the flagellum, and 11.1% (SD 7.7) with the parasite body. Conversely, an average of 41.4% (SD 7.2) *PFR2*^RNAi^ (+Tet) parasites were found to interact with the posterior; 49.5% (SD 5.8) with the flagellum, and 9.1% (SD 6.4) with the parasite body. No significant differences exist between lines. All data for Figure 6 is found in **Supp. Table 5**.

Having observed significant differences between the PFR2 knockdown and the parental and uninduced lines in terms of chip colonization and trap invasion (**Figure 5**), we went on to investigate whether differences existed in terms of parasite behaviour in the proximity of the traps. As previously described, parasites of the 2913 parental line predominantly interacted with the corner of the trap (58.7%), with 19.9% of interactions being unsuccessful, 18.1% contact elsewhere in the trap, and 3.3% enter and then exit (**Figure 6C**). In parasites of the *PFR2^RNAi^* (-Tet) line, the predominant behaviour was similar as in the parental line, namely, interaction with the corner of the trap (40.3%). However, a higher proportion of parasites compared to the parental line, failed to enter the trap (30.1%) or interacted elsewhere (26.2%). Only 3.5% of parasites entered the trap and then exited – a proportion similar to the one observed in the 2913 parental line (**Figure 6C**). In induced conditions, the predominant phenotype of *PFR2^RNAi^* cells was failed entry to the trap, with 59% of parasites displaying this behaviour. This is 2 to 2.6-fold higher than the control cell lines. Among parasites that did approach the trap, 24.5% persistently interacted with the corner, 16% made contact elsewhere in the trap, and only 0.5% entered the trap and then exited (**Figure 6C**).

To better understand the behaviour of parasites of all 3 samples, first we identified cells that could move freely within the confinement of the chip, to determine their mean square displacement over time. Free movement was mostly observed at the edges of the chip, in the region neighbouring the outer traps. The tracks of at least 60 cells per parasite line were obtained, and a significant defect in motility was confirmed in the *PFR2^RNAi^* cells where RNAi had been induced (blue) compared to the parental (black) and uninduced (red) lines (**Figure 6D**). Having identified that parasites swimming normally preferentially localize close to the corner of the V-shaped trap, we analysed the frequency of parasite positioning at various distances from the corner across all 3 parasite lines. We determined that while the 2913 and *PFR2*^RNAi^ (-Tet) lines preferentially localize within 0-20 µm (i.e one body length) from the corner, the induced counterparts show significantly smaller frequencies of localization in this proximity, with the majority of those entering the V-shaped trap localizing 30-60 µm away from the corner (**Figure 6E**).

In addition to the preferential localization at the V-shaped trap corner, we intriguingly observed that the direction of movement of the free-swimming *T. brucei* parasites was not always the usually observed flagellum-first direction. Rather, among 500 parasites observed (of the parental line), 40.3% were found interacting persistently with the trap corner, but performing ‘reverse’ swimming (yellow). Namely, these parasites were beating the flagellum, but cell body movement in the direction of the flagellum did not occur (i.e. the cell body was immotile) (**Video 7**). Instead, the parasite’s posterior was closest to the corner, without motion in the opposite direction (**Figure 6F and 6G**). In 48.6% of parasites, the conventional motion was observed, whereby the flagellum was leading, and was closest to the trap’s corner (black) (**Figure 6F and 6G**). Finally, in a minority of parasites (11.1%), the parasite’s body was bent and it was the parasite’s body, rather than the anterior or posterior ends, which constantly interacted with the trap’s corner (blue). Upon investigation of whether these types of interaction are conserved across parasite lines, we observed that this proportion is relatively similar and unaffected by the *PFR2*^RNAi^ knockdown (**Figure 6H**).

### Long-term confinement of trypanosomes in V-shaped traps

Next, we determined whether longer imaging times were possible. We began by determining which percentage of traps retained parasites for a minimum of 1 hour, and found that in 17% of traps, one or more trypanosomes remained in focus and within the confines of the V-shaped trap for 1 hour or more (**Figure 7A**). Upon comparing the 3 samples previously described, we found that although the proportion of traps invaded differ significantly between the parental, uninduced and induced lines, amongst those traps that were invaded, relatively equal proportions of traps had parasites that remained in focus and in the field of view for 1 hour or more (∼16-18%) (**Figure 7B, Video 5**).

**Figure 7.**
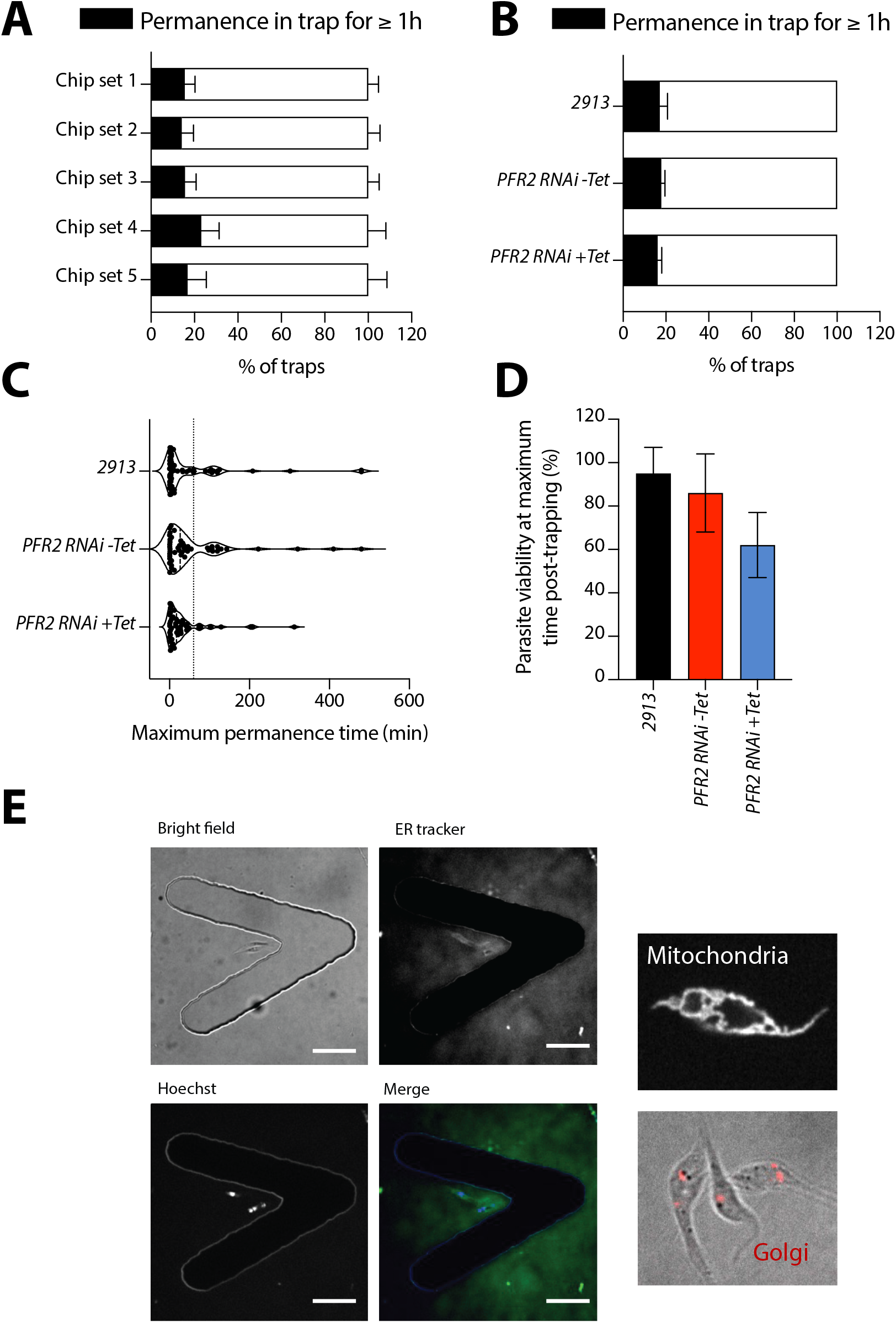
Tryp-Chip allows for long-term confinement of *T. brucei.* **A.** Investigation of *T. brucei* permanence within traps in Tryp-Chip allowed us to determine parasite relative immobilization for imaging for at least 1 hour. We found that on average, 17.08% of parasites imaged remained in place for >1 hour while 82.9% did not. **B.** Comparisons of parasite retention by parasite cell line showed that on average, 17.9% of *PFR2*^RNAi^ (-Tet) parasites could be imaged for > 1 hour, while 82.1% moved out of the field of view. Finally, 16.1% of *PFR2*^RNAi^ (+Tet) parasites could be imaged for >1 hour but 83.9% moved out of the field of view. **C.** While the prior information only sets 1 hour as a threshold, we also went on to determine the mean, median and maximum permanence time of all 3 parasite lines. The 2913 parental line had an average permanence time of 56.3 minutes, a median of 8, and a maximum of 480. The *PFR2*^RNAi^ (-Tet) had an average permanence time of 61.1 minutes, a median of 26.5 and a maximum of 480. Finally, the *PFR2*^RNAi^ (+Tet) line had an average permanence time of 38.1 minutes, a median of 17.5 and a maximum of 312. The median and maximum permanence times were significantly different between lines, while the differences in averages was not significant. **D.** We went on to explore parasite viability during maximum imaging times. The parental 2913 line showed 95% viability, while the *PFR2*^RNAi^ (-Tet) parasite line showed an 86% viability and the *PFR2*^RNAi^ (+Tet) parasite line showed 62% viability. **E.** Microscopy panels showing staining of sub-cellular structures. Top left panel shows a bright field image of a trap with *T. brucei*. The top right panel shows staining with ER tracker. The bottom left panel shows staining with Hoechst, and the bottom right panel shows the merged channels. The right-most panels show high-magnification images of staining with Mito-Tracker (mitochondria) and BODIPY ceramide (Golgi). All data for Figure 7 is found in **Supp. Table 6**.

We then extended the time frame and aimed at determining the maximum permanence time of each parasite line, considering also viability. Notably, parasites were kept at 26°C for the entire imaging procedure. However, in the current setup, media exchange was not possible. For this experiment, we inserted 10^4^ parasites/ml into the chips. We found that the mean permanence time for the 2913 parental line was 56 minutes, with a maximum of 480 minutes (8 hours). For the *PFR2*^RNAi^ line (-Tet), the mean permanence time was of 61 minutes, with a maximum of 480 minutes (8 hours). Finally, the *PFR2*^RNAi^ line (+Tet) had a mean permanence time of 38 minutes, with a maximum of 312 minutes (5.2 hours) (**Figure 7C**). Notably, parasite viability at 1-hour post-trapping was 97% in 2913 parasites, 85% in *PFR2^RNAi^* (-Tet) and 60% in *PFR2^RNAi^* (+Tet) (**Figure 7D**). Viability was assessed by motility and morphology, namely, dead parasites were fully immotile and showed signs of degradation including blebbing. Note that *PFR2^RNAi^* (+Tet) show decreased motility, but are not fully immotile when alive.

As a proof of concept, we determined whether the microfluidic device and the traps would allow longitudinal follow-up of sub-cellular structures. We stained 2913 parasites with ER Tracker, Mito Tracker or Bodipy-Ceramide, for ER, mitochondria, or Golgi labeling respectively, as well as Hoechst to visualize the nucleus and kinetoplast. We were able to observe the various structures in free-swimming parasites within the V-shaped traps (**Figure 7E**).

### Chips allow the study of *T. brucei* collective behaviour

Having initially observed that the majority of 6 and 8 µm-high traps harbour multiple parasites, even when the initial input is in low concentrations (i.e. 10^3^ parasites/µl), we hypothesized that this could reflect a form of collective migration which has been a key topic of interest for *T. brucei* [30–33]. We went on to investigate whether our prototype could be a useful tool to investigate these phenomena, especially as we observed that the majority of parasites cluster in the vicinity of the trap corner (similar to what we observed in single cells) (**Figure 8A, Videos 7-14**). We first established the proportion of successfully invaded traps that harboured single and multiple parasites (**Figure 8B**) at various concentrations. We confirmed that among all invaded traps in any one chip, 82% of traps in chips where 10^3^ parasites were inserted, had multiple parasites per trap, while only 18% had single parasites. In chips where 10^4^ or 10^5^ parasites were inserted, 76% of traps harboured multiple parasites, and in chips where 10^6^ parasites were inserted, up to 93% of traps harboured multiple parasites. In all cases, there were empty traps available for parasites to invade, yet parasites preferentially moved to locations where other parasites were already present. Next, we quantified the average number of cells per trap, based on initial inoculum (**Figures 8C-8D**). For the 2913 parental cell line, the average number of parasites per trap was 4.6 (SD 0.34), 7.3 (SD 0.32), 8.3 (SD 0.29) and 10.6 (SD 0.5) upon an inoculum of 10^3^-10^6^ parasites respectively. Notably, the flagellum has been defined as a key organelle for quorum sensing and collective motility. Whether PFR alterations influence collective motility is unknown. As proof of concept, we investigated in our design, whether the PFR2 knockdown would behave differently in terms of trap invasion, compared to the 2913 parental line. To test this, we first analysed the uninduced cell line *PFR2^RNAi^* (-Tet). The average number of parasites per trap was 4.5 (SD 0.1), 6.8 (SD 0.45), 6.7 (SD 0.5) and 9.5 (SD 0.3) upon an inoculum of 10^3^-10^6^ parasites respectively. These values were not significantly different to the parental 2913 line. However, in the *PFR2^RNAi^* (+Tet) line, the average number of parasites per trap was 4.3 (SD 0.2), 4.1 (SD 0.25), 4.7 (SD 0.27) and 4.4 (SD 0.2) upon an inoculum of 10^3^-10^6^ parasites respectively. This difference was significant (p < 0.001), compared to the parental and uninduced control lines.

**Figure 8.**
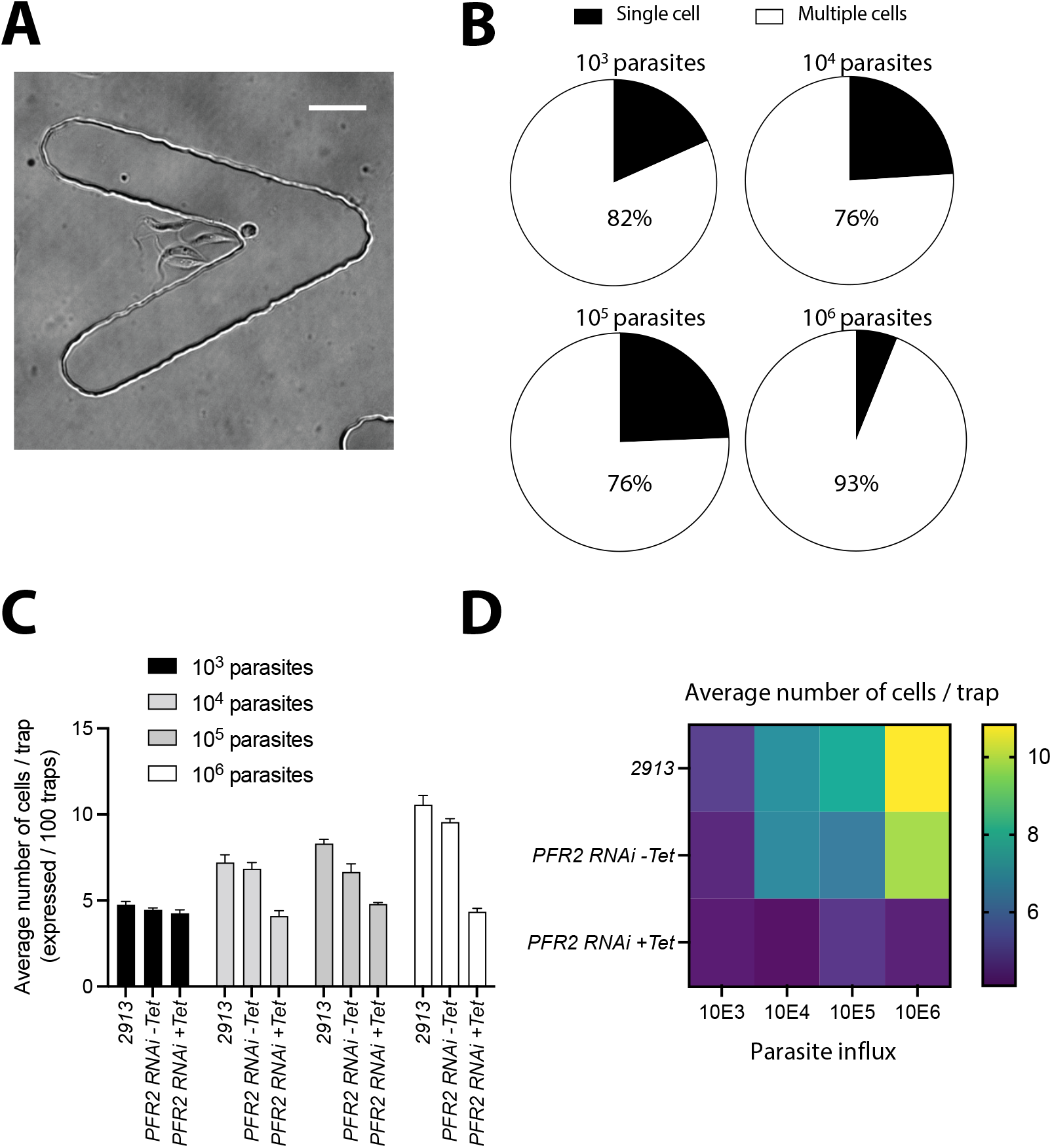
Tryp-Chip allows visualization of *T. brucei* collective behaviour. **A.** Bright field visualization allowed us to observe that most traps in Tryp-Chip are invaded by multiple parasites. **B.** 900 traps were measured, and values calculated from all traps invaded for over 1 minute. Invasion by multiple cells was the prevalent behaviour at 4 parasite concentrations tested. Influx of 10^3^ parasites resulted in 81.7% of invaded traps being invaded by multiple parasites, while 18.3% of those invaded, were invaded by single parasites. Influx of 10^4^ parasites resulted in 75.9% of invaded traps being invaded by multiple parasites, while 24.1% of those invaded, were invaded by single parasites. Influx of 10^5^ parasites resulted in 75.6% of invaded traps being invaded by multiple parasites, while 24.4% of those invaded, were invaded by single parasites. Influx of 10^6^ parasites resulted in 93.9% of invaded traps being invaded by multiple parasites, while 6.1% of those invaded, were invaded by single parasites. **C.** We went on to determine what the average number of parasites per trap was, depending on the parasite line at various influx concentrations. The average number of cells/trap when influx was 10^3^ parasites was 4.6 for the 2913 parental line, 4.5 for the *PFR2*^RNAi^ (-Tet) line, and 4.3 for the *PFR2*^RNAi^ (+Tet) line. The average number of cells/trap when influx was 10^4^ parasites was 7.3 for the 2913 parental line, 6.8 for the *PFR2*^RNAi^ (-Tet) line, and 4.1 for the *PFR2*^RNAi^ (+Tet) line. The average number of cells/trap when influx was 10^5^ parasites was 8.3 for the 2913 parental line, 6.7 for the *PFR2*^RNAi^ (-Tet) line, and 4.7 for the *PFR2*^RNAi^ (+Tet) line. The average number of cells/trap when influx was 10^6^ parasites was 10.6 for the 2913 parental line, 9.5 for the *PFR2*^RNAi^ (-Tet) line, and 4.4 for the *PFR2*^RNAi^ (+Tet) line. **D.** Heat map representation of average parasite number per trap for 3 parasite lines at 4 different influx parasite numbers. All data for Figure 8 is found in **Supp. Table 7**.

To further investigate whether this result was due to active parasite movement, rather than a flow-related artefact, we designed a setup whereby we inserted two sets of parasites. In the first part of the experiment, 10^4^ parasites were inserted via the inlet, and allowed to distribute and settle within the chip. Following 10 minutes, the entire chip was quantified, to determine the number of parasites per trap, as well as the number of traps that were occupied or empty. In the second part of the experiment, 10^4^ parasites were again inserted via the inlet, and allowed to distribute within the chip (which was already occupied by parasites). These were labelled with Mitotracker to discriminate them from the cells from the first influx (**Video 15**). Following 10 minutes, the entire chip was quantified, to determine the percentage of traps that were occupied or empty following the second parasite inflow (**Figure 9A**). Analysis of the 2913 parental line showed that on average, between 85 and 88% of new parasites (influx 2, labeled with Mito-Tracker) invaded occupied traps (unlabeled parasites), while only 12-15% went to empty ones (**Figure 9B**). This was consistent across two different parasite concentrations (10^4^ and 10^5^). A similar distribution was observed in uninduced *PFR2^RNAi^* (-Tet), whereby 77-82% of new parasites (influx 2) invaded occupied traps. This was also consistent across two different parasite concentrations (10^4^ and 10^5^) (**Figure 9B**). Conversely, the *PFR2^RNAi^* (+Tet) line showed a significantly different phenotype, with only 62-69% of parasites from influx 2 invading occupied traps (**Figure 9B**). To further explore potential passive and active trap invasion, we chemically fixed *T. brucei* parasites to investigate their distribution after influx into the chip both as a single influx, and upon double influx (as described in **Figure 9A**). While on average, 75% of traps were invaded by untreated 2913 parasites, only 16.2% of traps had parasites present if the parasites were previously fixed (**Figure 9C**). While in live cells invasion of traps was dynamic and continued over extended periods of time, fixed cell entry into traps did not vary after 30 seconds post-influx. Moreover, while an influx of 10^4^ live parasites resulted in an average of 7.8 parasites per invaded trap, influx of the same number of fixed parasites resulted in only 1.5 parasites per invaded trap. Upon investigating double influx (as described in **Figure 9A**), we found that if fixed cells were inserted into the device first, allowed to access traps, followed by insertion of live cells in a second influx, live cells showed no significant preference for empty or occupied traps (**Figure 9D**). This was independent of the parasite concentration inserted into the chip. Conversely, if live parasites were inserted first, and fixed parasites were inserted in a second influx and quantified in empty or occupied traps, only 15-20% of traps were ‘invaded’, regardless of whether these traps were empty or occupied during the first influx (**Figure 9E**). Findings in **Figures 9C-9E** are all consistent with a loss of capacity for active motion, and with a loss of capacity for chemotaxis – all consistent with fixation.

**Figure 9.**
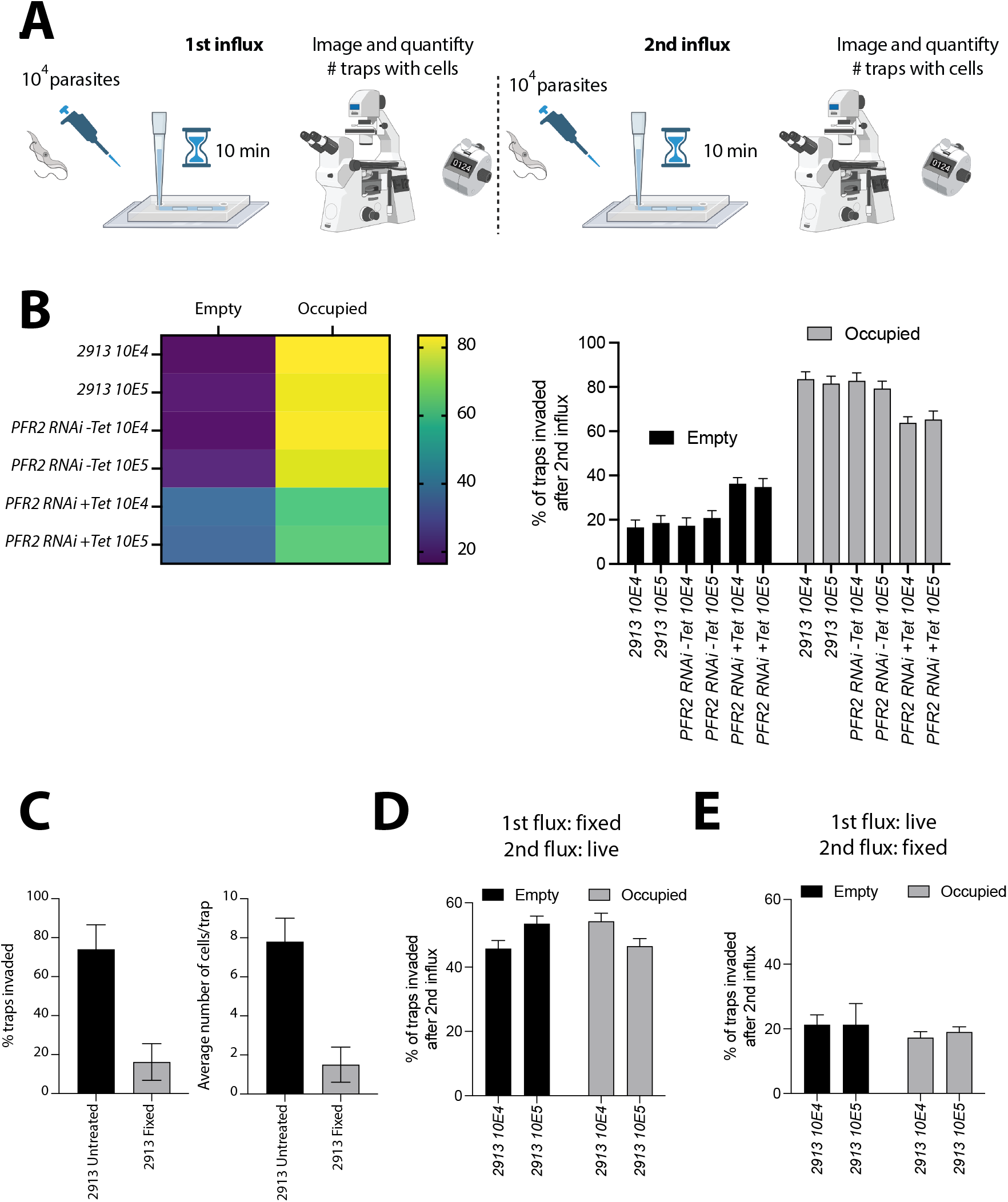
Tryp-Chip allows visualization of active *T. brucei* collective behaviour. **A.** Schematic representation of experimental design: 10^4^ parasites were inserted via the inlet, and allowed to distribute and settle within the chip. Following 10 minutes, the entire chip was quantified, to determine the number of parasites per trap, as well as the number of traps that were occupied or empty. In the second part of the experiment, 10^4^ parasites were again inserted via the inlet, and allowed to distribute within the chip (which was already occupied by parasites). These were labelled with Mito-Tracker to discriminate them from the cells from the first influx. Following 10 minutes, the entire chip was quantified, to determine the percentage of traps that were occupied or empty following the second parasite influx. **B.** The 2913 parental line showed that, on average, 85% of new parasites invaded occupied traps while only 15% invaded empty ones. This was consistent across two different parasite concentrations (10^4^ and 10^5^). In uninduced *PFR2^RNAi^* (- Tet), 82% of new parasites (influx 2) invaded occupied traps. This was consistent across two different parasite concentrations (10^4^ and 10^5^). The *PFR2^RNAi^* (+Tet) line showed a significantly different phenotype, with only 65% of parasites from influx 2 invading occupied traps. **C.** To explore potential passive and active trap invasion, we chemically fixed *T. brucei* parasites to investigate their distribution after influx into the chip both as a single influx, and upon double influx. While on average, 75% of traps were invaded by untreated 2913 parasites, only 16.2% of traps had parasites present if the parasites were previously fixed. Using live parasites, an influx of 10^4^ parasites resulted in an average of 7.8 parasites per invaded trap, while influx of the same number of fixed parasites resulted in 1.4 parasites per invaded trap. **D.** Using the double influx setup, we found that if fixed cells were inserted into the device first, allowed to access traps, followed by insertion of live cells in a second influx, live cells showed no significant preference for empty or occupied traps. All data for Figure 9 is found in **Supp. Table 8**.

## Discussion

### Technical features: strengths and future directions

We have presented here a microfluidic device with multiple V-shaped traps optimized for *T. brucei* capture and longitudinal visualization at both multiple and single cell level at high resolution, while allowing the parasite to remain free-swimming. This overcomes possible artefacts arising from immobilization by embedding in agarose, diminishing the temperature for imaging, or using chemicals such as poly-L-lysine for parasite adherence. The prototype we present here is complementary to various other valuable methods that have been generated over the last decade, to study various aspects of *T. brucei* behaviour [13,15,18,19,30–33] reviewed in [12]. We divide these methods into 6 groups for further discussion: a) temperature-based, b) chemical-adhesion-based, c) gel-based, d) optical traps; e) droplet-based and f) microfluidics and nanopatterning. Starting with the simplest methods, reducing the temperature to arrest motility is compatible with methods that require long acquisition times, or no motility in order to study protein-protein interactions. A limitation of this method, is that reduction of temperature could act as a confounder for the dynamic processes we wish to observe. As it reduces motility, it is incompatible with studies of native-state flagellar beating. Next, chemical immobilization methods (e.g. by the use of poly-L-lysine or concanavalin) are easy to use and in contrast to reduced temperature, allow for the study of protein dynamics (eg. by FRAP, FRET or photoconversion). However, it is based on the chemical adhesion of part of the *T. brucei* cell body and/or flagellum, which is a potential confounding factor for the study of dynamic processes. Equally, it is incompatible with the visualization of the free flagellum and imaging of free-swimming parasites. Agarose immobilization [37] and CyGEL immobilization [36] are also incompatible with visualization of the free flagellum. Optical trapping is a force nanoscopy-based method highly suitable for the study of propulsion forces generated by flagella [35]. While optimal for the study of single cells, it is not aimed at the study of collective behaviour. Emulsion droplets [18] are a relatively recent, valuable addition to the *T. brucei* toolkit. They are compatible with multiplex and large-scale screening, highly suitable for population studies. The method enables following up the progeny of single cells over various generations, and, importantly, it allows the study of free-swimming parasites. Its current design is not aimed at high temporal and spatial resolution, however, nor does it allow the study of parasite interactions with structures (simulating anatomical features).

Various microfluidic and micropatterning promising toolkits exist, each with its own strengths. One example is a large-scale screening microfluidic device generated by Hochstetter *et al* [34] which allows the study of parasite velocity and displacement. It currently does not aim at high temporal or spatial resolution, and is therefore incompatible with imaging sub-cellular dynamics. Equally it does not allow the study of parasite interactions with geometrical structures potentially mimicking anatomical features. Another device generated by Voyton *et al* [15] is also capable of multiplexing and large scale-screening, and enables perfusion to study parasite biochemistry. This design faces the same limitations as those described for the previously described microfluidic platform. Belonging to the group of nano- and micro-patterning are the designs generated by Heddergott *et al* [13] which are compatible with free-swimming parasites, with high-speed imaging and ideal for studies focusing on parasite navigation and interaction with obstacles. They are uniquely suited for 4D visualization. Their main limitation, however, is the scalability and ease of production and use. Our current work, presented here, aims to bring together several of the strengths of previously generated tools, to hopefully fill gaps in our capacity to address specific biological questions related to *T. brucei* biology.

Following various attempts at different trap geometries and measurements, the ones chosen for this work were the most promising ones, leading us to the prototyping of Tryp-Chip, presented in this work. V-shaped traps forming a tight angle at the corner (44°), and with arms of a length at least 2-fold the size of an average PCF *T. brucei* proved to be the most successful structures amongst those we tested. A key feature of this trap is the corner of the structure, which single parasites preferentially targeted, and into which multiple cells collectively accumulated (leaving other regions of the same trap relatively unoccupied). We found this observation interesting, given that it is similar to *T. brucei* behaviour previously described *in vivo* in the peritrophic matrix of its vector host, the tsetse fly [32,38]. While V-shapes are simple shapes, Tryp-Chip sets the basis for further investigations to replicate the geometry of the various environments that *T. brucei* finds in its insect and mammalian hosts. This would allow us to study the effect of environment geometry as one of multiple components governing parasite single and collective behaviour. Equally, a limitation of our current work is that we studied parasite behaviour without flow (other than the initial influx of parasites). The relevance of flow differs, depending on the parasite stage studies – for instance, while flow might be less relevant for procyclic stages once they have crossed the peritrophic membrane within the insect vector, it is of vital importance in bloodstream forms in blood and lymph. Since Tryp-Chip’s design is custom made, including inlet and outlets, flow can be controlled and modulated to mimic biologically relevant conditions. Implementing continuous flow could also allow us to study parasite motility (single and collective) in the presence of exogenous substances including trypanocidal drugs, host metabolites, and chemo-attractants, to name a few. Equally, due to the static nature of the chip in its current form, media exchange was not possible, making extended imaging over several days impossible due to desiccation unless the chip is sealed, lack of nutrients without media exchange, and/or parasite death. The implementation of flow would allow for this limitation to be overcome. Nevertheless, the current design allowed us to image parasites for extended periods of time already.

### Tryp-Chip as a tool to study free-swimming single cells

In this first prototype of Tryp-Chip, we explored the role of parasite motility in chip colonization and the potential for Tryp-Chip as a tool for single cell visualization, analysing the parasite’s permanence time in each trap. One of our main findings was that upon disruption of the *T. brucei* paraflagellar rod (by silencing PFR2), Tryp-Chip colonization was greatly altered compared to the results found in the control parasite population. Loss of PFR2 impacted time to full chip colonization, percentage of traps invaded, and overall displacement within the chip. The *PFR2*^RNAi^ knockdown was used as a proof of concept for the potential to study cell and flagellar motility. Previous work on PFR alterations has demonstrated an essential role for the PFR in flagellum and cell motility in *T. brucei* [20,22,39]. This proof-of-concept opens the door to the applicability of Tryp-Chip to other studies. A large body of work has demonstrated that the flagellum plays critical roles beyond parasite motility, including cell cycle progression, organelle segregation or immune evasion [4,31,34,42–58, 59]. Combined with novel tools such as the recent genome-wide subcellular protein map for *T. brucei* [60] and TrypTag [61,62] - a project that resulted in over 5000 fluorescent reporter lines – Tryp-Chip offers the possibility of phenotypic characterization of flagellar beating/cell movement, and its relation to cell cycle progression. Moreover, while our current work focused on PCFs, we envisage that Tryp-Chip could be a valuable tool to explore *T. brucei* heterogeneity. Specifically, our knowledge on mammalian tissue forms and ‘intermediate’ forms arising during the *T. brucei* life cycle remains limited [63]. Isolation of parasites from different host reservoirs ([32,38,64–73]), for their longitudinal observation in Tryp-Chip for characterization of multiple features, is an important future step we envisage. Given its potential to visualize single cell motility in free-swimming parasites, Tryp-Chip could be extended for use with other *Trypanosoma* species, and even other kinetoplastid parasites.

### Tryp-Chip as a tool to study collective behaviour

An interesting feature that Tryp-Chip allowed us to investigate, is the preferential accumulation of multiple parasites in single traps, despite space and traps not being restrictive. One common aspect to most motile organisms is their capacity to move in response to changes in the environment. Chemotaxis [74], social motility [reviewed in 4,79–82], and quorum sensing [77–79] are some of the responses displayed by microorganisms. Chemotaxis is a strategy by which an organism moves towards or away from a chemical cue, as an evolutionary response for protection, nutrient acquisition, and migration, including movement through physical barriers [discussed in 78]. In the context of *T. brucei*, social motility has been a topic of great interest for over a decade, with findings showing that *T. brucei* assemble into groups that engage in collective motility. While no exogenous chemical cues were introduced into Tryp-Chip, our results from double influx of cells showed that parasites were more likely to move into traps which already contained multiple parasites. While one hypothesis for this observation is that flows generated by the flagellar beating of the parasites already present in the traps results in preferential cell movement into occupied traps, our work using fixed cells suggests that parasite accumulation within single traps is not solely the result of flagellar beating-induced micro-currents. Another hypothesis is that parasites attract each other – this could be explained by mechano-sensing whereby live parasites are able to respond to changes in flow, or chemical cues, whereby groups of parasites signal their presence to others via secreted molecules. Furthermore, the accumulation of parasites in single traps was less pronounced in the *PFR2^RNAi^* mutants. This could be related to altered sensing and altered motility, but remains to be tested. While the relatively short time lapse used with this Tryp-Chip prototype does not allow us to reach conclusions regarding social motility (which requires several days), other studies have implicated signaling at the parasite’s flagellum in social motility responses [80–85]. Our findings on the *PFR2^RNAi^* mutants are consistent with the hypothesis that flagellar structural alterations result in reduced chemotaxis as well as reduced motility. While further work is required to disentangle the multiple possible components governing collective parasite behaviour in *T. brucei*, we believe that Tryp-Chip could be a valuable tool to screen social motility-altered mutants at single cell and collective levels. Moreover, Tryp-Chip could enable visualization of free-swimming parasites, and their response to parasite communities and geometric structures mimicking anatomical features of the insect and mammalian host.

Examples of questions we could address with Tryp-Chip include: How do different parasite stages (or even parasite species) differ in terms of cell cycle progression and motility? Or how do they respond to different geometrical structures (mimicking anatomical features) in Tryp-Chip? Altogether, the prototype we present here, has multiple potential applications to answer fundamental questions on *T. brucei* biology, including the link between flagellar structure and function, and collective parasite behaviour.

## Acknowledgements

The authors thank Parul Sharma, Charlotte Izabelle, Christelle Travaillé and Audrey Salles for helpful input. We thank the Bastin and Glover groups for helpful discussions. MDN was funded by an HFSP postdoctoral fellowship (LT000047/2019-L). We acknowledge Claudio Franco (Instituto de Medicina Molecular, Lisbon, Portugal) for key intellectual input. Work in the Trypanosome Cell Biology Unit (Bastin Lab) is funded by an ANR grant (ANR-18-CE13-0014-01) and by a French Government Investissement d’Avenir programme, Laboratoire d’Excellence “Integrative Biology of Emerging Infectious Diseases” (ANR-10-LABX-62-IBEID). We gratefully acknowledge the UTechS Photonic BioImaging (Imagopole), C2RT, Institut Pasteur, supported by the French National Research Agency (France BioImaging; ANR-10–INBS–04; Investments for the Future).

## Author contributions

Conceptualization, M.D.N., S.G.; methodology, M.D.N., S.G.; software, M.D.N., E.F., S.G.; validation, M.D.N. and S.G.; formal analysis M.D.N., P.B., S.G.; investigation M.D.N., E.F., S.G., P.B.; data curation, M.D.N.; visualization, M.D.N., S.G.; writing – original draft, M.D.N., S.G., P.B.; writing – review & editing, all authors.; supervision, S.G., P.B.; project administration, M.D.N., S.G., P.B..; funding acquisition, M.D.N., S.G., P.B.

